# Notch-1 mediates Epithelial-Mucosal Healing during Murine Colitis Recovery Phase

**DOI:** 10.1101/2022.06.13.496012

**Authors:** Latika Luthra, Sahil R. Patel, Zhen Bian, Yuan Liu, Didier Merlin, Pallavi Garg

## Abstract

Inflammatory bowel disease (IBD) is marked by inflammation mediated epithelial-mucosal damage. The intestinal epithelial forms a tight barrier displaying two contrasting functions: restricting the entry of potentially harmful substances while, on the other hand allowing the selective passage of nutrients. The damaged epithelial-mucosal barrier causes the exposure of mucosa layers to luminal inflammatory contents. This eventually leads to the leaky epithelium, exposing the immune cells, release of various cytokines, and results in loss of epithelial homeostasis. Therefore, maintenance of healthy epithelial-mucosal lining is critical during IBD recovery. Notch signaling is an evolutionarily conserved molecular pathway crucial for the development and homeostasis of most tissues. Notably, the deregulation of Notch signaling is involved in IBD (Ulcerative colitis and Crohn’s disease). Notch signaling also plays a vital role in wound healing and tissue repair. Here, we investigated the role of Notch-1 signaling in wound healing and regeneration of colonic epithelium during colitis recovery phase by using conditional deletion of Notch-1 in colonic epithelium of mice. We used colonic carcinoma cell line HCT116 transiently transfected with Notch1 intracellular domain (NICD) to support in vivo data. We observed that deletion of Notch1 among mice was associated with compromised healing after colitis. Therefore, targeting the Notch-1 pathway might provide a novel therapeutic strategy for the patients recovering from colitis.

## INTRODUCTION

The luminal surface of the colon is continuously exposed to a plethora of foreign antigens, including food, foodborne pathogens, and commensal bacteria. A single layer of columnar epithelial cells covers the intestinal mucosa and serves as a barrier to protect the host from invasion by potential pathogens. Various types of epithelial cells forming a crypt in the mucosa perform diverse functions. Epithelial stem cells at the crypt bottom continuously divide to produce transit-amplifying cells, which can then divide several times before creating terminally differentiated epithelial cell lineages: absorptive enterocytes and three types of secretory epithelial cells, namely, goblet cells, enteroendocrine cells, and Paneth cells.

Under physiological conditions, the cellular composition of the intestinal epithelium keeps renewing itself and establishing homeostasis; however, disruption of this homeostatic state has been associated with various gastrointestinal diseases, including ulcerative colitis (UC) [1]. Ionizing radiation causes severe damage to healthy tissue. The gastrointestinal tract is a radiation-sensitive organ, and we used this strategy to induce colitis in mice. Notch-1 proteins function as receptors for transmembrane ligands, Jagged and Delta-like proteins, to regulate a broad spectrum of cell fate decisions. Notch-1 receptors undergo γ-secretase–mediated proteolytic cleavage upon ligand activation to release the Notch intracellular domain (NICD). The liberated NICD translocate into the nucleus, where it forms a transcriptional activator complex with other transcriptional factors to regulate the Notch signaling pathway [2].

However, the involvement of Notch-1 in the pathophysiological changes occurring in irradiated colonic tissue is still unknown. To better understand the role of Notch-1 signaling in epithelial defense functions and gut immune homeostasis, we have generated mice in which there is an intestinal epithelial cell (IEC)-specific deletion of the Notch-1 gene (Notch-1^ΔIEC/ ΔIEC^) and exposed them to ionizing (X ray) radiation. Irradiation induced DNA damage in colonic epithelial cells leading to the loss of tight junctions *(Claudin, Occludin*, and *Zonula occludens* (*ZO1*), which are essential for epithelial integrity. A leaky epithelium is permeable to all sorts of pathogens, causing inflammation. Previous studies with transgenic expression indicated that NICD inhibits secretory cell differentiation with a reciprocal increase in immature progenitors [2, 3]. These observations support the biological significance of Notch signaling not only in the binary cell fate decision and in the maintenance of the proliferating progenitors in the crypts of the intestinal epithelium. So far, the contribution of Notch signaling to epithelial defense functions remains elusive. It is known that Notch signaling in intestinal epithelium is upregulated in the inflamed mucosa of UC patients [4]. We are directing toward the crucial role of Notch in assuring epithelial integrity and rapid turnover of the cells to close the wound/ulcers created during ablation by inflammation mediated injury.

## MATERIALS and METHODS

### Animal Models

Mice carrying a floxed Notch-1 allele Notch-1^Flox/Flox^ (using the symbol Notch-1^F/F^) were obtained from the Jackson Laboratory. To generate Notch-1^ΔIEC/ ΔIEC^ (using the symbol Notch-1^ΔIEC^) mice, we crossed Notch-1^F/F^ mice with Villin-Cre transgenic mice obtained from The Jackson Laboratory. Notch-1^ΔIEC^ mice and control Notch-1^F/F^ littermates were maintained under specific pathogen-free (SPF) All animal procedures were performed in accordance with the Guide for the Care of Use of Laboratory Animals. Eight-weeks old, both female and male, Notch-1^F/F^ (wild type) and Notch-1^ΔIEC^ mice were used. Mice were genotyped by PCR analysis as previously described. The Notch-l suppression among Notch-1^ΔIEC^ mice was identified by using PCR genotyping using Notch forward primer (5’-TGCCCTTTCCTTAAAAGTGG-3’) and reverse primer (5’-GCCTACTCCGACACCCAATA-3’).

### DSS Acute Colitis Induction and Recovery Phase

Eight weeks-old, both female and male, Notch-1^F/F^and Notch-1^ΔIEC^mice were administered 1.5% Dextran sodium sulfate (DSS) (MP Biomedicals, Solon, OH) through their drinking water for 5 days to induce acute colitis. Mice were then left to recover on water (no DSS added) for 7 days. Mice were euthanized on day 12. Control group mice, Notch-1^F/F^and Notch-1^ΔIEC^, were given regular water for the same amount of time. Body weight and stool consistency for all the mice were recorded during both, DSS and recovery phase.

### Irradiation Injury by X-rays

Eight weeks-old, both female and male, Notch-1^F/F^and Notch-1^ΔIEC^mice were anesthetized with an intraperitoneal injection of 80 mg/kg ketamine and 10mg/kg xylazine. Mice whole body irradiation was performed with a single exposure to 2Gy abdominal irradiation at a dose rate of 1.2 Gy/min using a Rad Source RS 2000 Small Animal Irradiator (thanks to Zhen Bian-Post doc from Dr Liu’s lab). Intestinal samples were harvested 7 days later for histologic evaluation. Body weight for all the mice were recorded during before irradiation and subsequently for a week. On day 7 animals including control mice which were not exposed to X-rays were sacrificed and their colon was removed.

### Bacterial Culture and Plasmid Preparation

DH5α Competent Cells were mixed with pCMV-IRES2-EGFP plasmids (Epoch Life sciences) for expressing a gene together with EGFP. Vectors with or without the hNICD1 gene (2412 bp) between Nhe1 HF and EcoR1 HF restriction sites and transformed through heat shock. Cells were incubated on ice for 20 minutes, spread on Nutrient broth (BD Biosciences) agar plates, and cultured in an incubator overnight. Single colonies were moved to liquid broth and cultured for 8 hours in a shaking incubator. 200µl of the broth was moved to 500ml liquid broth (Sigma Aldrich) culture flasks and incubated overnight in a shaking incubator. Cells were isolated by centrifuging cultured broth, and plasmids were prepared using (Invitrogen, PureLink™ HiPure Plasmid Midiprep Kit), following the manufacturer’s protocol. The obtained plasmid DNA was resolved in nuclease/endotoxin-free water and stored at -20°C.

### Cell Culture, Transfection, Wound Healing Assay, and Migration Assay

HCT116 cells, colon human carcinoma cells, were cultured McCoy’s 5a Medium Modified (ATCC, catalog No. 30-2007) supplemented with 10% FBS (Atlanta Biologicals), and 100 units/ml penicillin, and 100 mg/ml streptomycin (GIBCO) at 37°C in 5% CO_2_. Cells (5 × 105) were seeded in 6 well plates and incubated for 24 hrs. When cells were 70-90% confluent, cells were then transfected with hNICD1 cloned into pCMV-IRES2-EGFP vector or without hNICD1 (Empty vector) pCMV-IRES2-EGFP. The plasmids were transiently transfected into HCT116 cell line using Lipofectamine 3000 transfection kit (Invitrogen) according to the manufacturer’s protocol. 24 h after transfection, monolayer was wounded by scratching method with a sterile pipette tip and images were taken at 24, and 48 hr by using Keyence BZ-X700 microscope. Transfection efficiency was done by looking at the change in enhanced green fluorescent protein (EGFP) signals in fluorescence microscope.

For migration-24 hr after transfection, cells were treated with mitomycin C (10 µg/ml) for 3 hours and washed 3 times with PBS to inhibit cell proliferation. Scratch assay was performed to assess cell migration. The pictures were taken at time points 0, 24, or 48 observing the width of the cell free gap closing. Experiments were performed in duplicate and on at least on two separate occasions. Cell migration was assessed by measuring the remaining open area of the wound by ImageJ software.

### Protein Extraction and Western Blot (WB) Analysis

Colonic epithelial mucosal stripping was obtained from whole colon of Notch-1^F/F^ (wild type) and Notch-1^ΔIEC^ (conditional knockout) mice with and without exposure to X ray irradaition. Protein for WB was extracted from the stripping using Pierce RIPA buffer (Cat# 89900, Thermo Fisher Scientific) mixed with Halt Protease & Phosphatase Inhibitor Cocktail (100×, Cat# 78440, Thermo Fisher Scientific). Samples were quantified using the DC protein assay (Bio-Rad). Quantified samples were prepared with LDS sample buffer with or without β-mercaptoethanol (Sigma-Aldrich) for nonreducing gels. Prepared samples were boiled at 58°C for 10 minutes and briefly centrifuged. 10% Criterion™ TGX™ Precast Midi Protein Gel, 18 well were loaded with protein samples, β-tubulin (Sigma-Aldrich) and GAPDH (Abcam)was used as loading control. Followed by electrophoresis in Tris-glycine SDS buffer. Proteins were transferred to nitrocellulose membranes that were then blocked with TBS-T supplemented with 5% skimmed milk or 5% filtered-BSA. Primary antibodies ATOH1 (BD Biosciences), Hes-1 (Affinity Bioreagents), Claudin 2 (Invitrogen), Claudin 1 (Invitrogen), NF-kB (Invitrogen), Occludin (Invitrogen), GAPDH (cell Signaling), ZO1 (Cell Signaling) were diluted in 5% skimmed milk or 5% filtered-BSA and incubated overnight at 4°C with rocking. Membranes were washed and incubated with horseradish peroxidase (HRP)-conjugated secondary antibodies (goat anti-mouse secondary antibody (Bio-Rad) or goat anti-rabbit secondary antibody (Abcam)) diluted in 5% skimmed milk for 3 hours. Chemiluminescence was optimized by ECL (Super signal West Pico Plus, Thermo Fisher Scientific), and signals were pictured using BIO-RAD CHEMIDOC touch imaging system.

### Mouse Colonoscopy

Notch-1^F/F^ and Notch-1^ΔIEC^ mice with and without radiation exposure were subjected to colonoscopy to assess the mucosal layer thickness, signs of inflammation, and dysplastic lesions. This was performed using the colonoscope (Xenon Nova 47S, STORZ).

### Hematoxylin and Eosin (H&E) Staining and Histological Score Evaluation

Formalin fixed and paraffin embedded colon Swiss rolls from Notch-1^F/F^ and Notch-1^ΔIEC^ mice with and without radiation exposure were stained with H&E. Histological scores were evaluated based on inflammation infiltration of white blood cells, crypt damage, and foci of ulceration in the entire colon. Images were taken using Keyence BZ-X700 microscope at X10, X20 and X40 magnification.

### Alcian Blue-Periodic Acid Schiff (AB-PAS) Staining

Formalin fixed and paraffin embedded colon Swiss rolls from Notch-1^F/F^ and Notch-1^ΔIEC^ mice with and without radiation were used. Sections were deparaffinized, hydrated and incubated with acetic acid for 3min. Sections were stained with 3% Alcian Blue (Sigma-Aldrich). Next, sections were stained with periodic Acid Solution for 5 minutes at room temperature (18–26°C). After washing, sections were incubated with Schiff’s Reagent for 15 minutes at room temperature (18–26°C). Rinsed the slides again and counterstain slides in Hematoxylin Solution, Gill No. 3, for 90 seconds. The slides were washed, and sections were then hydrated with alcohol, cleared with xylene, and sealed. Images were taken using Keyence BZ-X700 microscope at X20 magnification.

### RNA Extraction and QPCR Analysis

Total RNA was extracted from colonic tissues using the RNeasy mini-Kit (Qiagen) according to the manufacturer’s instructions. Then, complementary DNA was generated from the total RNA isolated using the Maxima first-strand complementary DNA synthesis kit (Thermo Scientific). The cDNA samples were amplified with the DreamTaq Green PCR Master Mix (2X) (Thermo Scientific) and the primer sets specific for mouse genes. The sequences of the primer sets are shown in the table. Target gene expression was assessed by a comparative cycling threshold method, using expression of 36BR gene levels as the internal standard. Realtime QPCR data is depicted as fold change (DDCT values). Bacterial genomic DNA was isolated by gently scraping the inside of the colon. Bacterial genomic DNA samples were amplified with a Eppendorf (Realplex 4 Mastercycler epgradient S 5345 Real Time PCR 96 Well) using the DreamTaq Green PCR Master Mix (2X) and the primer sets specific for bacterial 16S rDNA.

**Table 1.**
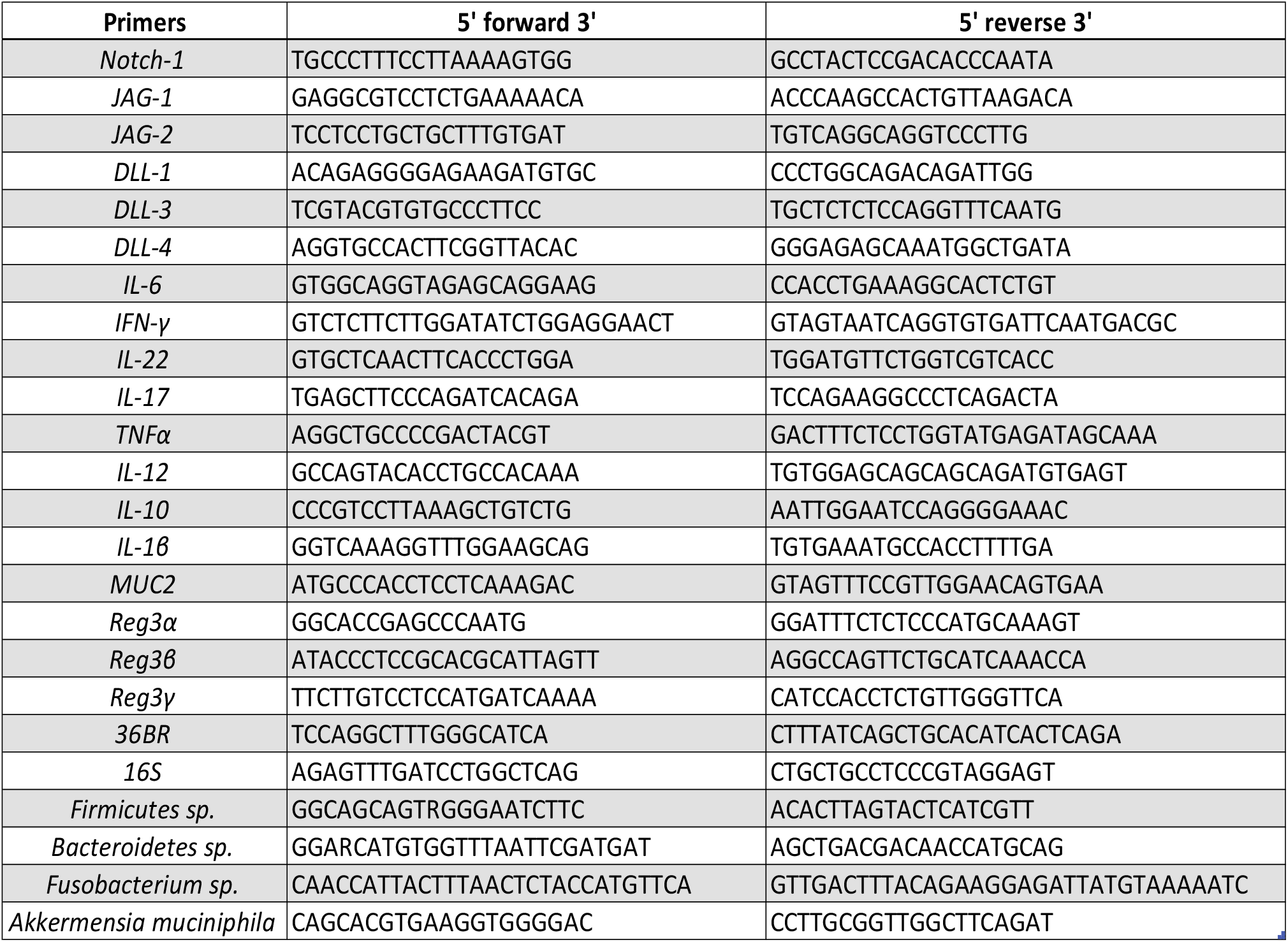
Sequences of the QPCR primers used in the study.

### Immunofluorescence Staining of Colon Swiss Rolls

Notch-1^F/F^ and Notch-1^ΔIEC^ mice with and without radiation colon Swiss rolls were formalin fixed and paraffin embedded for immunofluorescence staining. 5-μm-thick paraffin-embedded sections of colon tissue samples were deparaffinized by xylene and dehydrated by different concentration of alcohol solutions and then were treated with citrate buffer (pH 6.0) for antigen retrieval. Immunostaining was performed according to manufacturer’s protocol using the Tyramide SuperBoost™ Kits with Alexa Fluor™ Tyramides (Invitrogen). 3 % H_2_O_2_ was used to block endogenous peroxidase for 15 min, followed by diluted goat serum to reduce nonspecific staining for 30 min. After that, the sections were incubated with primary Ki67 (1:200, ab16667, Abcam), MUC2 (GeneTex), Lgr5 (Santa Cruz Biotechnology), ZO-1 (Cell signaling), Cyclin D1(A-12, Santa Cruz Biotechnology), Lysozyme C (E-5, Santa Cruz Biotechnology) at 4 °C overnight. Washed the slides three times with PBS and added 2–3 drops (approximately 100–150 µL) of anti-mouse/anti rabbit poly-HRP-conjugated secondary antibody to the tissue and incubate for 60 minutes at room temperature. Rinsed the section for 10 minutes with PBS at room temperature. Applied 100 μL of the Alexa Fluor™ 594/488 Tyramide reagent and incubated for 2–10 minutes at room temperature. Reaction was ceased using stop reagent. Counterstain the section with Phalloidin iFluor 488/555 diluted with 1X PBS for 1 h. Finally, the sections are mounted with ProLong Gold antifade reagent with DAPI (Invitrogen) with coverslip. Images were taken with Keyence BZ-X700 microscope.

### Apoptotic Cell Labeling via TUNEL Staining

Paraffin sections of colons were deparaffinized and apoptotic cells were identified by immunofluorescent terminal deoxynucleotidyl transferase-mediated deoxyuridine triphosphate nick-end labeling (TUNEL) staining using In Situ Cell Death Detection Kit, Fluorescein (Millipore-Sigma) according to the manufacturer’s instructions. Quantification of apoptosis was performed by counting the number of apoptotic cells in a crypt and was shown as percentage values.

### Statistical Analysis

Differences between two groups were analyzed by Student t test. When variances were not homogeneous, the data were analyzed by Mann Whitney U test. Differences among more than two groups were analyzed by one-way ANOVA followed by Dunnett test. When variances were not homogeneous, the data were analyzed by Kruskal–Wallis test. All data was expressed as means± SEM. P values <0.05 was considered statistically significant [ns = non-significant, ** = p<0.01, *** = p<0.001, **** = p<0.0001]. Statistical tests were applied using GraphPad Prism (version 6).

## RESULTS

### Conditional Deletion of Notch-1 in Colonic Epithelium Compromises the Architecture and Cell Differentiation in the Colon

To directly assess the role of epithelial Notch signaling in intestinal homeostasis, we generated Notch-1^ΔIEC^ mice. These mice were born at the expected Mendelian ratio. The Notch l deletion among Notch-1^ΔIEC^ mice was identified by using PCR genotyping. (Fig 1A) shows the genotyping result where lanes 1 and 2 are homozygous Notch-1^ΔIEC^ mice expressing only one band (281 bp) while lanes 4 and 5 are heterozygous mice, expressing two bands, the mutant and the Notch-1^F/F^ (281 and 231 bp, respectively). (Fig 1B) To confirm Notch-1 presence at the protein level, protein lysates were prepared from Notch-1^F/F^ mice vital organs such as the colon, small intestine, liver, kidney, lung, and stomach. All the organs respectively indicate the presence of Notch-1, depicting that Notch-1 is essential in all organs to maintain homeostasis. Histological analysis of Notch-1^ΔIEC^ mice showed a massive cellular infiltration, disrupted crypt architecture, increased neutrophil infiltration, and a few foci of ulceration compared to Notch-1^F/F^ mice (Fig 1C and 6D). The histological score is calculated using three parameters described in the methods section. The bar graph shows that Notch-1^ΔIEC^ mice had a significantly higher histological score (5.5 ± 0.5) compared to Notch-1^F/F^ mice (1.5 ± 0.5). Notch-1 mediates the enterocyte population against goblet cells and mucus secretory cells during stem cell differentiation in the colon. AB-PAS staining of these 8-wk-old Notch-1^ΔIEC^ mice demonstrated that the number of secretory cell lineages was increased in the colon (Fig 1E), an observation consistent with a previous observation in mice with inducible Notch-1 deficiency [5]. The acidic mucins are identified by Alcian blue and neutral mucins by PAS; wherever crypt mixtures occur, the resultant color will depend upon the dominant moiety. Notch signaling drives early crypt progenitors to control their lineage determination. Phenotypic alterations occurred mainly in the crypts and were homogeneous throughout the intestinal and colonic mucosa in Notch-1^F/F^ mice (Fig 1E), demonstrating highly efficient excision of the Notch-1-floxed alleles by the Cre transgene and the absence of a mosaic phenotype. Goblet cell hyperplasia was particularly obvious in the colon of Notch-1^ΔIEC^ mice. AB-PAS images indicated that Notch-1^ΔIEC^ had more goblet cells compared to Notch-1^F/F^ mice. (Fig 1F) is the bar graph representation of the goblet cell hyperplasia due to the conditional deletion of Notch-1 signaling. It shows quantification of goblet cells, indicating that Notch-1^ΔIEC^ mice had a significantly increased no. of goblet cells (22.3 ± 0.3) compared to Notch-1^F/F^ mice (6.2 ± 0.7). This data indicates that Notch-1^ΔIEC^ mice have a defect not only in the regulation of IEC differentiation but also in the maintenance of gut homeostasis.

**Fig 1.**
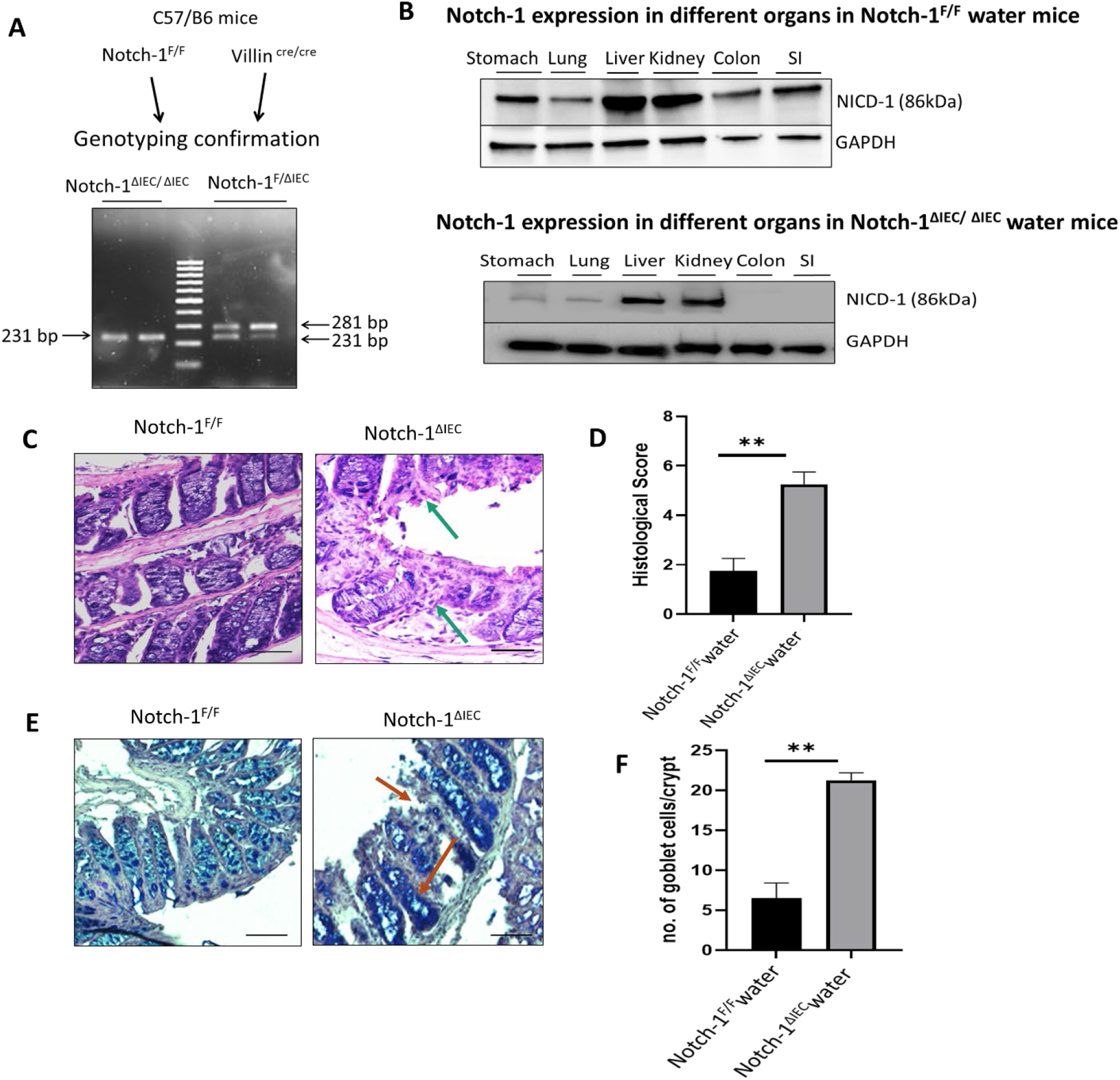
Conditional deletion on Notch-1 signaling in mice colon leads to the loss of epithelial crypt architecture and increased goblet cell population. Notch-1^ΔIEC^ mice were generated using the breeding scheme for generating Cre/*lox* tissue-specific knockouts by The Jackson Laboratory Cre/*lox* system. Notch-1^F/F^ mice were used as a control. (A) Qualitative PCR of the DNA extracted from mice tails with the primers designed by the supplier (Jackson Laboratories). Homozygous Notch-1^ΔIEC^ mice have one mutant band at 281 bp, while heterozygous mice have two bands, the mutant at 281 and the Notch-1^F/F^ at 231 bp. Lane 3 is a 100 bp ladder. Each lane shows blots performed loading (25μg/lane) using whole organ cell lysate of the colon from Notch-1^ΔIEC^ and Notch-1^F/F^ groups. (B) Expression of NICD-1 in different organs in homeostasis is probed using an anti-NICD antibody, and GAPDH was used as a loading control. (C) Colonic tissue sections were stained with H&E for histological examination in the presence and absence of Notch-1. (D) The histological colitis score was calculated based on the criteria described in the materials and methods section. Scale bars, 100 μm. Data are representative of three independent experiments. (E) AB-PAS staining for detection of goblet cells (lower panels). (F) The number of goblet cells per crypt was quantified. Values are mean ± SD (n = 5). **p < 0.01 (Mann–Whitney U test).

### Notch-1 Signaling Mediates Inflammation in DSS-Induced Acute Colitis

Eight weeks old, both female and male, Notch-1^F/F^ and Notch-1^ΔIEC^ mice were given 1.5% DSS to induce acute colitis. Body weight of mice was monitored throughout the experiment and mice were euthanized at day 12. (Fig 2A) shows the schematic of DSS acute colitis recovery model. (Fig 2B) Shows the bodyweight change in percentage indicating a significant increase in body weight among Notch-1^F/F^ mice compared to Notch-1^ΔIEC^ mice during the recovery period. (Fig 2C) To assess inflammation among Notch-1^F/F^ and Notch-1^ΔIEC^ mice in acute colitis, H&E staining was performed. H&E images indicate crypt architecture damage, infiltration of neutrophils, and loci of ulceration among Notch-1^ΔIEC^ mice compared to Notch-1^F/F^ mice in acute colitis. (Fig 2D) Bar graph representation of histological score calculated on three parameters: infiltration of neutrophils, loss of crypt structure, and foci of ulceration. (Fig 2E) shows colonoscopy images indicating the thinning of mucosa and bleeding among Notch-1^ΔIEC^ mice compared to Notch-1^F/F^ mice. This data confirms that Notch-1^ΔIEC^ mice are taking longer to recover from DSS induced colitis in comparison to Notch-1^F/F^ mice. This observation might suggest that Notch-1 is essential for faster healing.

**Fig 2.**
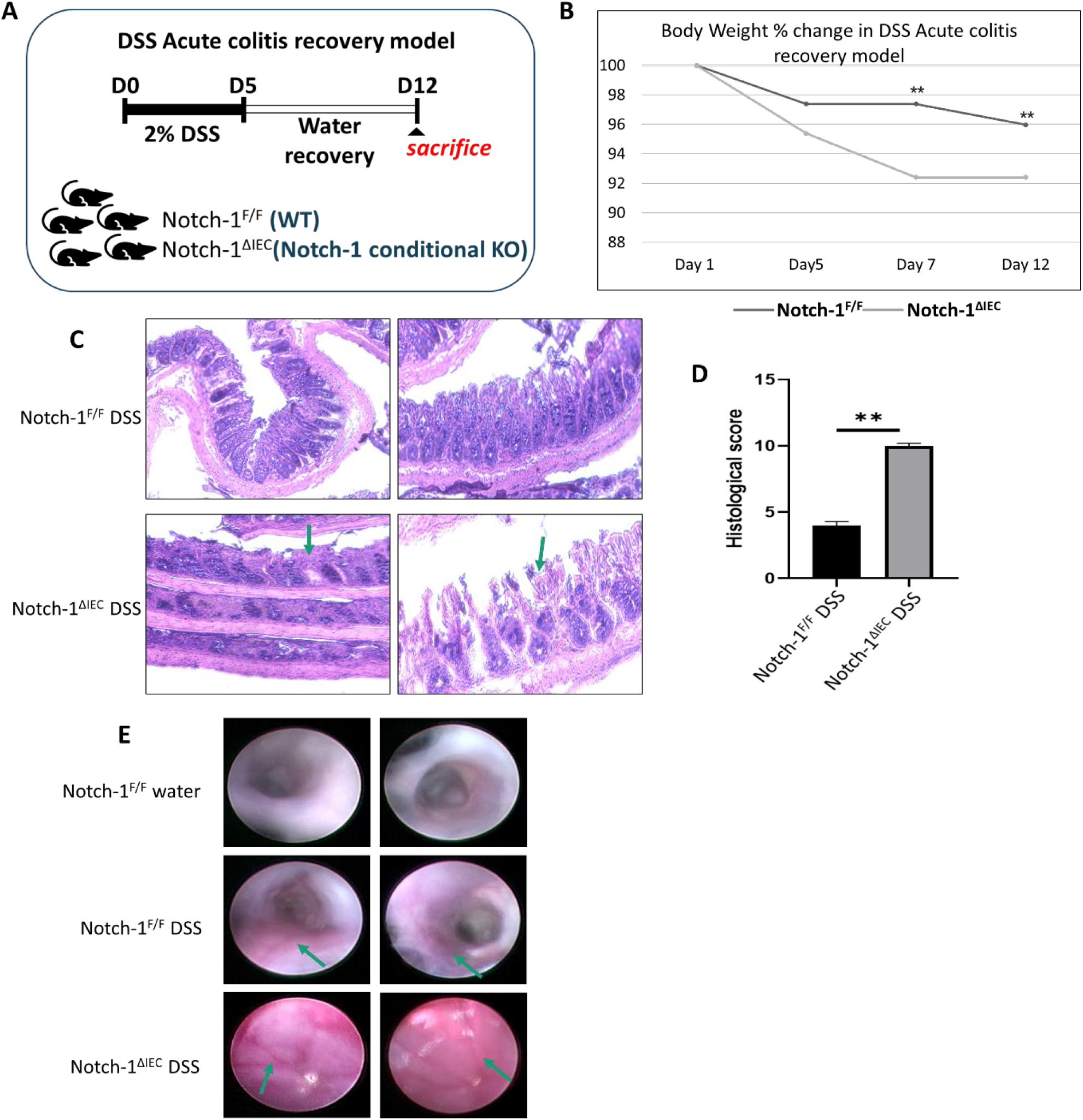
Notch-1 signaling mediates inflammation in DSS-acute colitis mice. DSS induced damage in the colon of Notch-1^F/F^ and Notch-1^ΔIEC^ mice (A) The schematic of DSS model of inducing colitis in Notch-1^F/F^ and Notch-1^ΔIEC^ mice (n=5/group) and recovery for seven days. (B) Changes in body weight were monitored throughout the experiment, and mice were euthanized on day 12. (C) HE staining of colon swiss rolls from Notch-1^F/F^ and Notch-1^ΔIEC^ mice given 1.5% DSS in water. Scale bars are 100µm (D) Histology score based on HE staining method. Data were expressed as mean ± SD (Student t-test). * p < 0.05, ** p < 0.01 vs. 1.5% DSS groups (Notch-1^F/F^ and Notch-1^ΔIEC^). (E) Colonoscopy images to show inflammation in colon increased slightly in Notch-1^F/F^ mice after DSS induced colitis but Notch-1^ΔIEC^ mice had escalated inflammatory response. Scale bars, 100 μm.

### Notch-1 Affects Production of Proinflammatory Cytokines in DSS Colitis in Mice

To assess the inflammatory environment, mRNA levels of cytokines that are known to mediate acute inflammatory progression in experimental colitis were measured by qPCR as described in the methods section. Interestingly, the mRNA levels of all pro-inflammatory cytokines were higher in Notch-1^ΔIEC^ mice as compared to Notch-1^F/F^ mice (Fig 3A). These results together with the inflammation observed via H and E staining and colonoscopy indicate that Notch-1 signaling mediates DSS-induced colitis and is associated with upregulation of pro-inflammatory.

**Fig 3.**
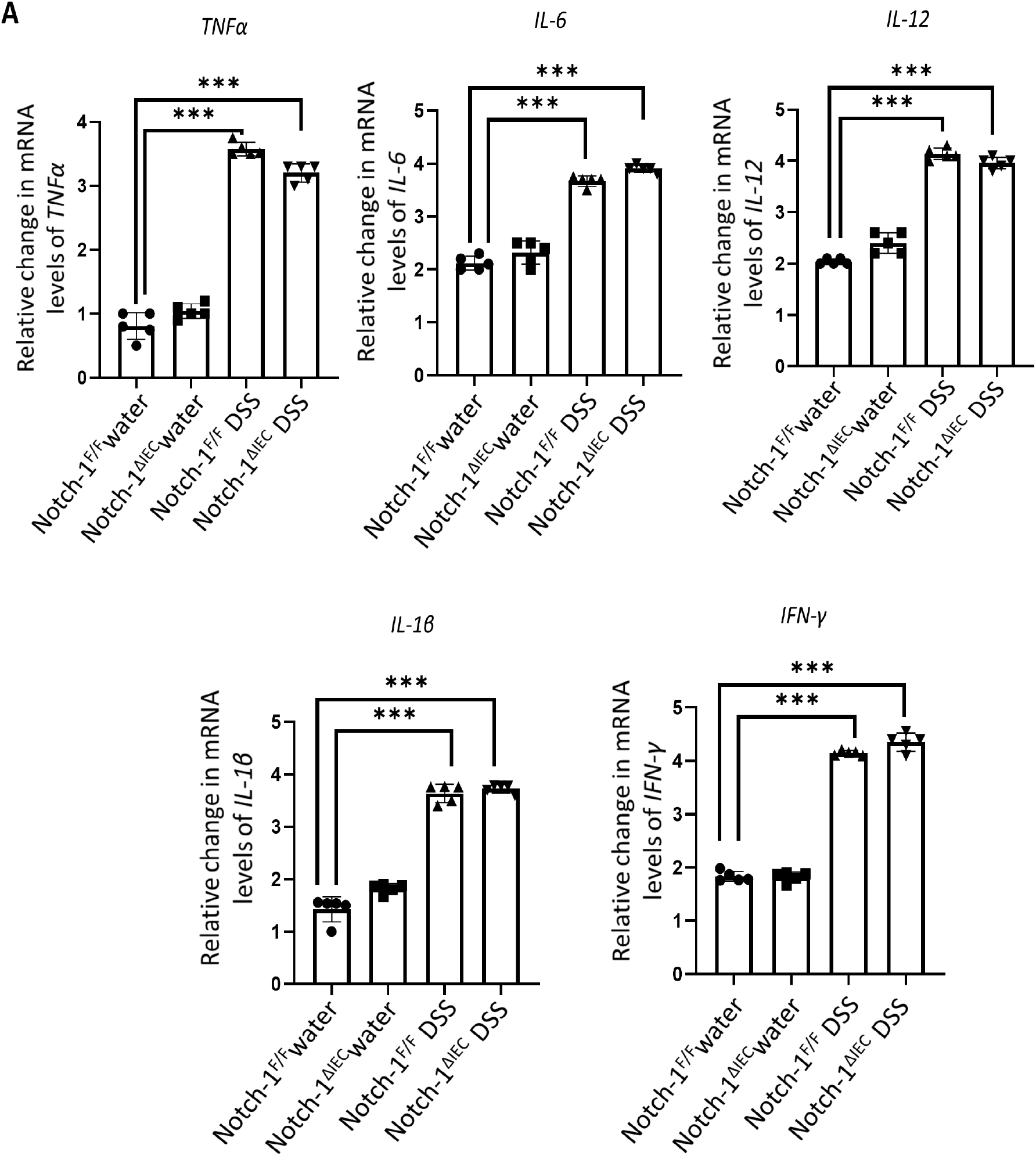
Notch-1 affects production of proinflammatory cytokines in DSS colitis in mice. (A) mRNA expression of cytokines including TNF, IL-6, IL-12, IL-1β and IFNγ was examined by qRT-PCR in colonic mucosa from 8-week-old mice after 12-day recovery from DSS acute colitis. Data is depicted as fold change. Data were normalized to expression of 36BR (housekeeping gene). Data were expressed as mean ± SD (n=5) of three independent experiments. * p < 0.05, ** p < 0.01, *** p < 0.001, **** p < 0.0001, vs. radiation groups (one-way ANOVA test followed by Dunnett test).

### Deletion of Notch-1 in IECs Induces Significant Epithelial Damage During Colitis

Eight weeks old, female and male, Notch-1^F/F^and Notch-1^ΔIEC^mice, were irradiated with a single exposure to 2Gy abdominal irradiation at a dose rate of 1.2 Gy/min small animal irradiator (Fig 4A). Intestinal samples were harvested seven days later for histologic evaluation. Body weight for all the mice was recorded before irradiation and subsequently for a week. (Fig 4B) During the experiment, the body weight of animals in the Notch-1^ΔIEC^ irradiated group continued to decrease significantly from day 3. Compared with the Notch-1^F/F^ group, body weight loss dramatically improved due to the lack of the Notch-1 gene. (p < 0.01). (Fig 4C) In addition, a western blot was run to check the protein levels of Notch-1 in different groups, clearly depicting increased Notch-1 levels in Notch-1^F/F^ as compared to Notch-1^ΔIEC^, which doesn’t have the gene, but Notch-1^F/F^ Notch-1 levels increased in colitis over Notch-1^F/F^ control mice never treated with radiation [6]. In line with the above findings, epithelial crypt damage, mucosa edema, depletion of goblet cells, and inflammatory cell infiltration was observed in HE staining of the, Notch-1^F/F^and Notch-1^ΔIEC^mice, but more damage is observed in Notch-1^ΔIEC^mice as compared to, Notch-1^F/F^mice (Fig 4D).

**Fig 4.**
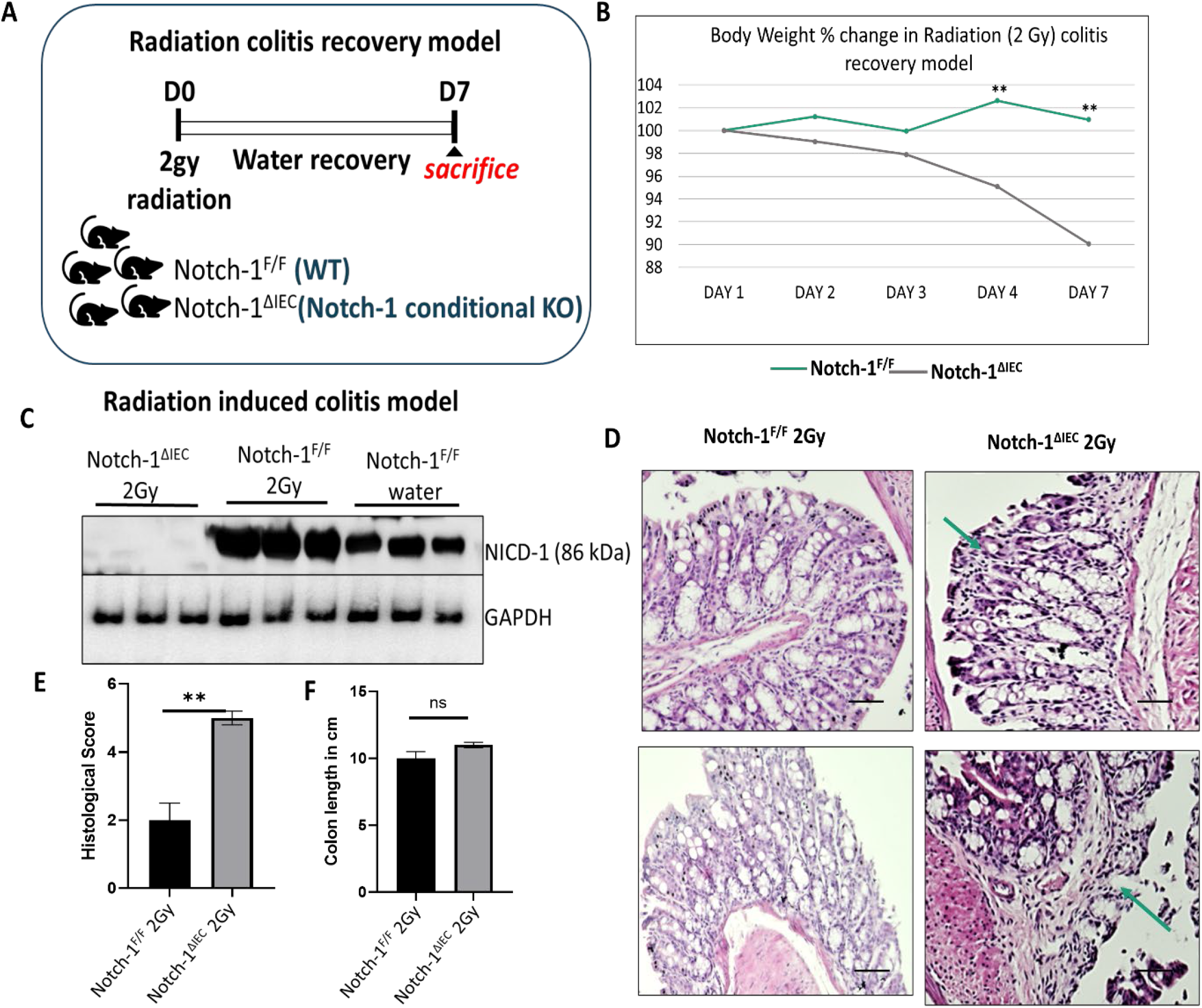
Notch-1 relieves radiation-induced colitis in mice. Radiation-induced damage in the colon of Notch-1^F/F^ and Notch-1^ΔIEC^ mice (A) The schematic of radiation model inducing colitis in Notch-1^F/F^ and Notch-1^ΔIEC^ mice (n=5/group) and recovery for seven days. (B) Changes in body weight were monitored throughout the experiment, and mice were euthanized on day 7. (C) Western blot comparing Notch-1 expression in radiated mice (Notch-1^F/F^ and Notch-1^ΔIEC^) vs. control. (D) H&E staining of colon swiss rolls from Notch-1^F/F^ and Notch-1^ΔIEC^ mice exposed to radiation (magnification X10, X20, and X40). Scale bars are 100µm (E) Histology score based on H&E staining method. (F) Statistics of colon length. Data were expressed as mean ± SD (Student t-test). * p < 0.05, ** p < 0.01 vs. radiation groups (Notch-1^F/F^ and Notch-1^ΔIEC^).

Furthermore, the histology score, assessing the severity of colitis, increased distinctly after radiation treatment, whereas it was markedly lower in the Notch-1^ΔIEC^ as compared to the Notch-1^F/F^ group. Nevertheless, the Notch-1^F/F^ group exhibited obvious protection from mucosa damage and less histological inflammation, which showed a lower histopathological score (Fig 4E). Although there were clinical manifestations of bloody diarrhea and body weight reduction, the colon length was unchanged in Notch-1^F/F^ and Notch-1^ΔIEC^ groups before and after radiation (Fig 4F). Combined, these results demonstrate that Notch-1 exerts therapeutic effects on radiation-induced colitis.

### Notch-1 Ameliorates Radiation-Induced Epithelial Permeability by Enhancing the Function of Tight Junctions

Depending on the nature and the site of the inflammation, occludin, ZO-1, and different claudin proteins have been implicated in the pathogenesis of IBD [7]. Increased expression of claudin-1, -2, and -18 [8-12] and downregulation of claudin-3, -4, and -7 were reported in ulcerative colitis [9, 10]. Replacement of barrier-forming claudins with pore/channel-forming claudins like claudin-2 influences ion and fluid movement across cells, which is reflected in disease symptoms, including diarrhea. Analysis of many tight junctions in the presence or absence of Notch-1 in colitis showed that the lack of a Notch-1 in Notch-1^ΔIEC^ mice dramatically increased claudin-2 expression. Conversely, occludin expression was markedly decreased. The effect on claudin-1 was relatively mild, although it was also reduced (Fig 5B). ZO connects junctional proteins such as occludin and claudin to the actin cytoskeleton, and these protein interactions maintain TJ formation and function. There was a marked reduction in ZO-1 expression in Notch-1^ΔIEC^ mice exposed to radiation colitis (Fig 5A). To define the mechanism that associates Notch-1 signaling and barrier function and inflammatory mediators influence on transcriptional regulation of claudins in the TJ, we performed western blotting of NF-κB and STAT3 (Fig 5C). Pro-inflammatory cytokines such as TNF-α, IL-1β, and IFNγ promote TJ permeability. TNF-α suppresses TJ barrier function due to the activation of the NF-κB pathway and decreased ZO-1 protein level. IL-1β increased TJ permeability via activation of the NF-κB pathway[13]. (Fig 5D) To test the role of cytokines in inflammation and how they affect different TJ proteins, the levels of TNF-α, IL-1β, and IFNγ were checked via QPCR. These data were correlated by immunofluorescence staining (Fig 5E). We confirmed that claudin-2 was localized to the cell junction and was upregulated when Notch-1 was knocked out. In the physiologic state, claudin-2 expression is restricted to proliferative colonic crypt base epithelial cells. During mucosal inflammation, claudin-2 expression is upregulated in cells, and its expression extends beyond the crypt-base proliferative cells in the colon[14]. Moreover, this is depicted in the immunofluorescence images crypt to upper compartment movement of claudin-2. Therefore, the absence of Notch-1 mediated disruption of this stoichiometry may account for the abnormal barrier function. Altogether, these findings provide evidence for the involvement of the Notch-1 signaling pathway not only in barrier function but also in the architecture of the epithelium.

**Fig 5.**
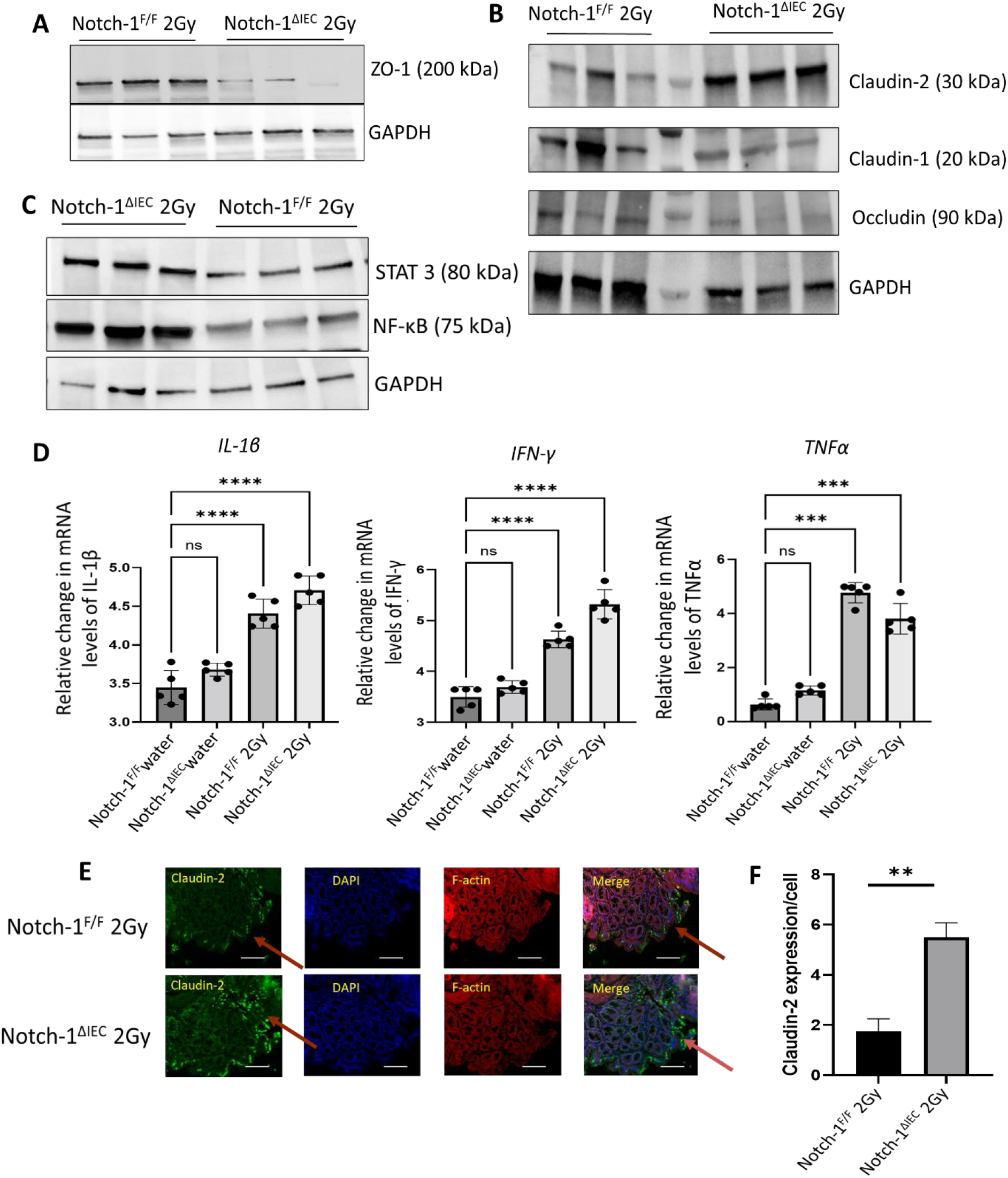
Notch-1 ameliorates radiation-induced epithelial permeability by enhancing the function of tight junctions. (A) and (B) Immunoblotting for claudin 2, claudin 1, occludin, and ZO-1 in membrane protein extracts obtained from the colon of radiation-induced colitis mice groups (Notch-1^ΔIEC^ and Notch-1^F/F^). (C) Expression of inflammatory mediators in colitis in Notch-1^ΔIEC^ and Notch-1^F/F^ mice. (D) qRT-PCR was used to examine the mRNA expression of TNF-α, IL-1β, and IFN-γ. Data were expressed as mean ± SD of three independent experiments. (0ne-way ANOVA) * p < 0.05, ** p < 0.01, *** p < 0.001, **** p < 0.0001, vs. radiation groups (Notch-1^ΔIEC^ and Notch-1^F/F^). (E) Immunofluorescence staining for Claudin-2 TJ in the distal colon of 8 wk-old Notch-1^ΔIEC^ mice and control Notch-1^F/F^ (2Gy and water group) using mAbs against Cld-2 (green), nuclei were counterstained with DAPI (blue). Scale bars, 100 μm. (F)Quantification of Claudin 2 expression/cell in 2Gy condition in Notch-1^ΔIEC^ and Notch-1^F/F^ mice. Data are representative of three independent experiments. Values are mean ± SD (n = 5). *p < 0.05 (Student t-test).

### Notch-1 Inhibits Apoptosis and Promotes Proliferation in the Colonic Epithelium in Mice with Radiation-Induced Colitis

The disruption of homeostasis between proliferation and apoptosis in the colonic crypts is engaged in the pathogenesis of UC[15-17]. So, the proliferation and apoptosis of colon epithelial cells were examined by immunostaining of Ki67 to test cell proliferation and apoptosis by TUNEL assay. Ki67+ proliferating cells were abundant at the bottom of virtually all the crypts in control mice, whereas Ki67+ cells were nearly absent in multiple crypts of Notch-1^ΔIEC^ mice (conditions-water or radiation-induced colitis) (Fig 6A). Notch-mediated Hes1 expression contributes to the cell proliferation in the intestinal crypts by transcriptional repression of the Cyclin D1 regulator [18]. Immunostaining analysis detected Cyclin D1+ cells in the bottom of every crypt in control mice; however, such cells were nearly absent in the crypts of Notch-1^ΔIEC^ mice (Fig 6B). Western blot of Hes1 protein suggests that reduction in epithelial cell proliferation may result from the downregulation of Hes1 because of the absence of Notch-1 (Fig 6C). These data also raise the possibility that Notch-1^ΔIEC^ mice are defective in epithelial cell turnover and activate apoptosis. We, therefore, performed TUNEL assay (Fig 6D).

**Fig 6.**
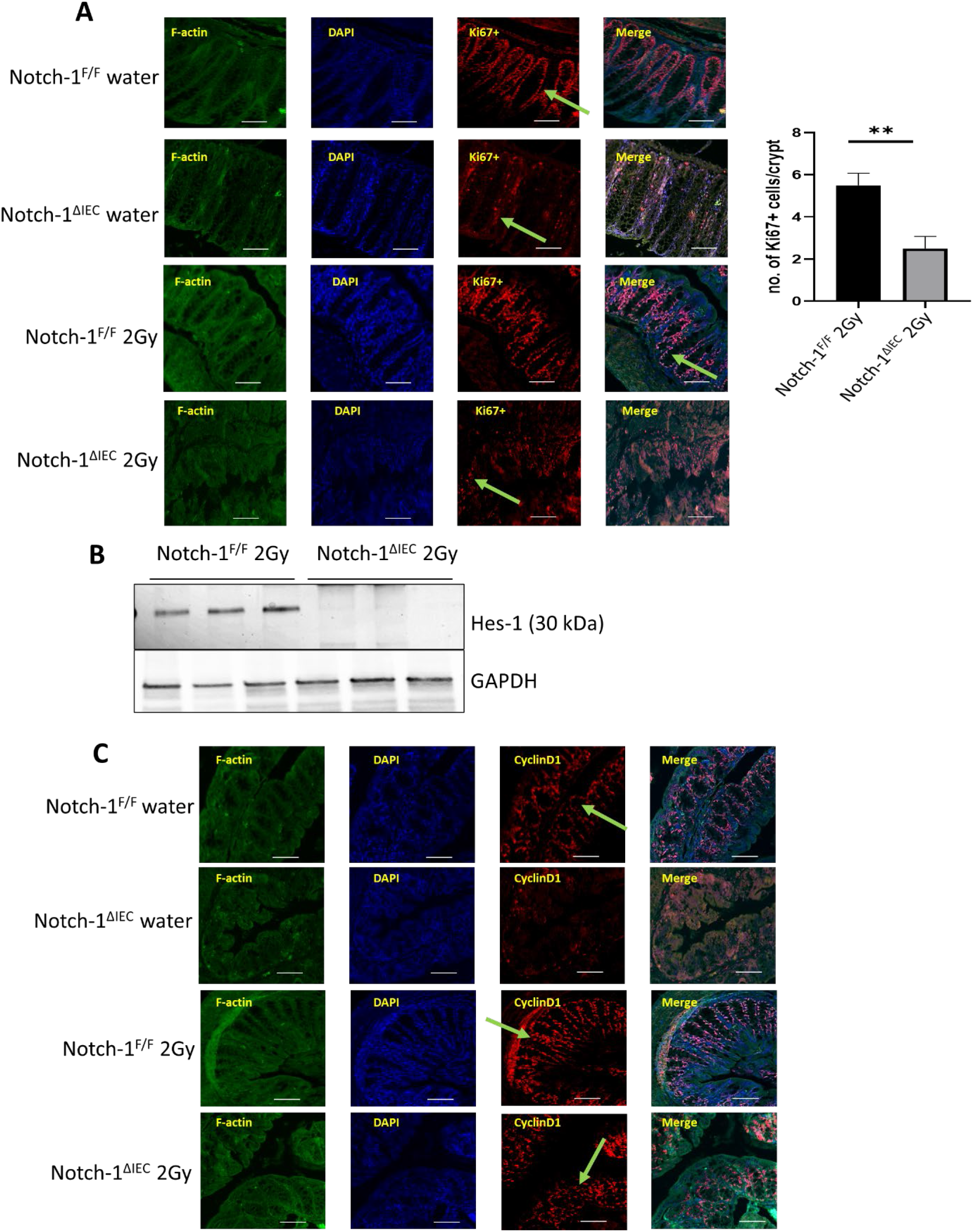
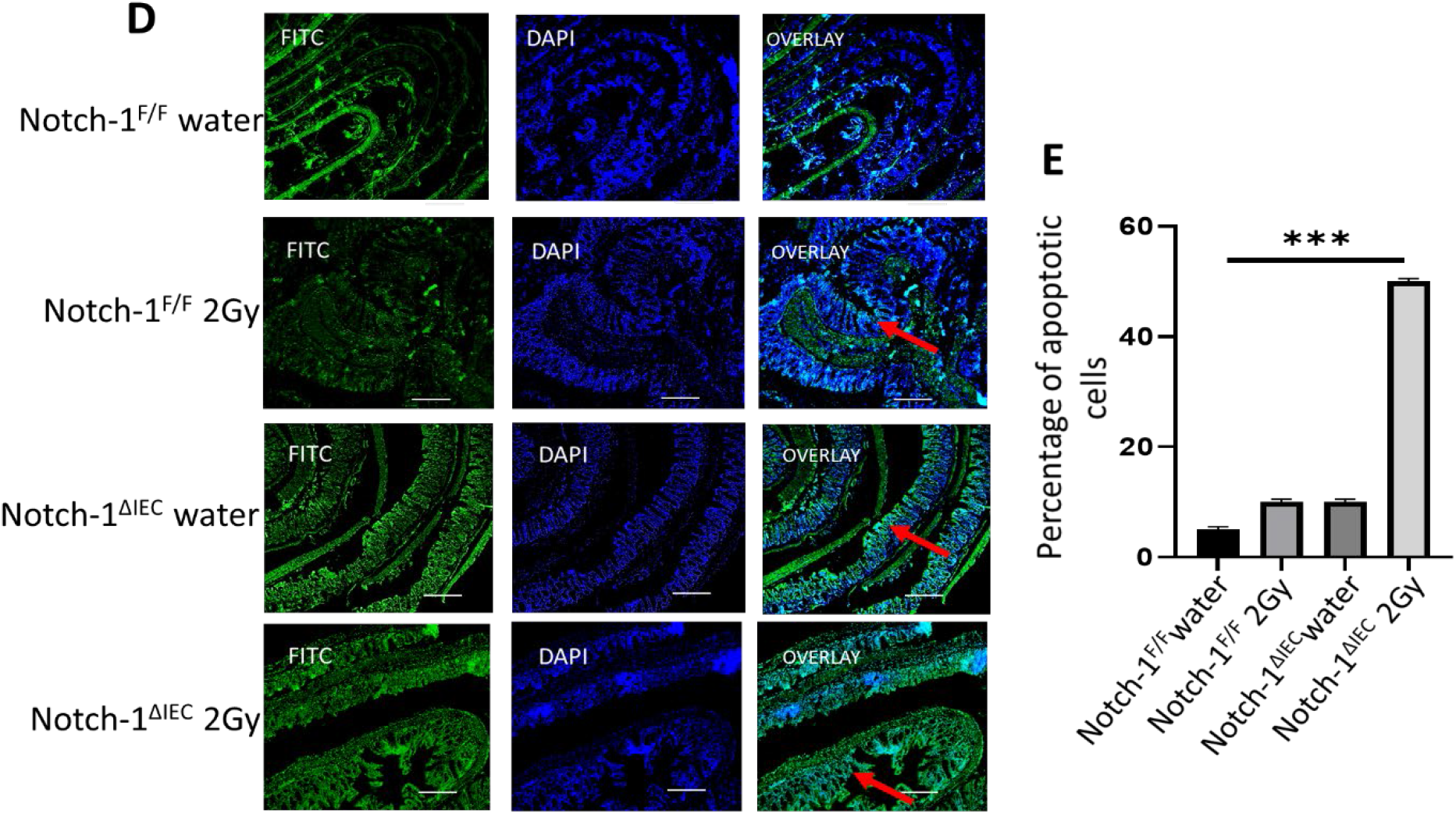
Notch-1ΔIEC mice have a reduction in apoptotic cells and increment in proliferating epithelial progenitors in the colon. (A) Immunofluorescence staining of the distal colon from Notch-1^ΔIEC^ mice and control Notch-1^F/F^ littermates was performed using polyclonal Ab against Ki67. The number of Ki67+ proliferating cells per crypts was quantified. Values are mean ± SD (n = 5). **p < 0.01 (Mann–Whitney U test). Scale bars, 100 μm. (B) Immunoblotting of the distal colon from radiated Notch-1^ΔIEC^ mice and control Notch-1^F/F^ littermates was performed using Ab against Hes1. Data are representative of three independent experiments. (C) Immunofluorescence staining of the distal colon from Notch-1^ΔIEC^ mice and control Notch-1^F/F^ littermates was performed using Ab against Cyclin D1. (D) Epithelial cell apoptosis in Notch-1^ΔIEC^ mice and Notch-1^F/F^ mice. TUNEL assay (In Situ Cell Death Detection Kit) was used to visualize apoptotic cells with fluorescein-dUTP (green), followed by DAPI for nuclei counterstaining (blue). Scale bars, 100 μm. (E) For quantification analysis, 20 crypts per mouse were randomly selected, and the number of green cells per crypt was counted. ***p < 0.001 (Mann–Whitney U test).

The hallmark of apoptosis is DNA degradation. The DNA cleavage may yield DNA breaks (nicks) which can be detected by TUNEL (**T**dT-mediated d**U**TP-X **n**ick **e**nd **l**abeling). (Fig 11E) To quantify the rate of colonic epithelial apoptosis, we counted the number of apoptotic cells per crypt. Consistent with Ki67-staining data, the number of apoptotic cells was significantly increased in the colonic epithelium of Notch-1^ΔIEC^. Overall, these results suggest that Notch-1 helps maintain the intestinal epithelium barrier by preventing apoptosis and enhancing proliferation.

### Notch Enhances Intestinal Barrier Function and Epithelial Cell Differentiation of Radiation-Induced Colitis Mice

Mucus secreted from the crypt mixes with Paneth cell secretions containing antibacterial peptides, lysozyme, DMBT1, and MUC2 [19-23]. The Paneth cell products will, together with enterocyte-produced antibacterial proteins like Reg3γ, generate an antibacterial gradient in the mucus and keep the bacteria away from the epithelial cell surfaces. Notch signaling in the colon is implicated in the maintenance of stem cells and progenitor cells and the inhibition of goblet cell differentiation. In the intestine, Notch signaling, especially Notch1, regulates stem cell-fate determination in the crypt [24]. Once stimulated, activation of Notch upon ligand binding promotes the expression of Hes-1 (Hairy and Enhancer-of-split-1), which then would suppress downstream ATOH1, resulting in an increase of enterocytes in parallel with restricting differentiation into goblet cells and MUC2 expression (Fig 7A). Enhanced levels of MUC2 are observed in Notch-1 conditional knockouts. [25-27]. Moreover, Notch-1^ΔIEC^ mice exhibited goblet cell hypertrophy and hyperplasia as identified by Alcian Blue (Fig 7B)

**Fig 7.**
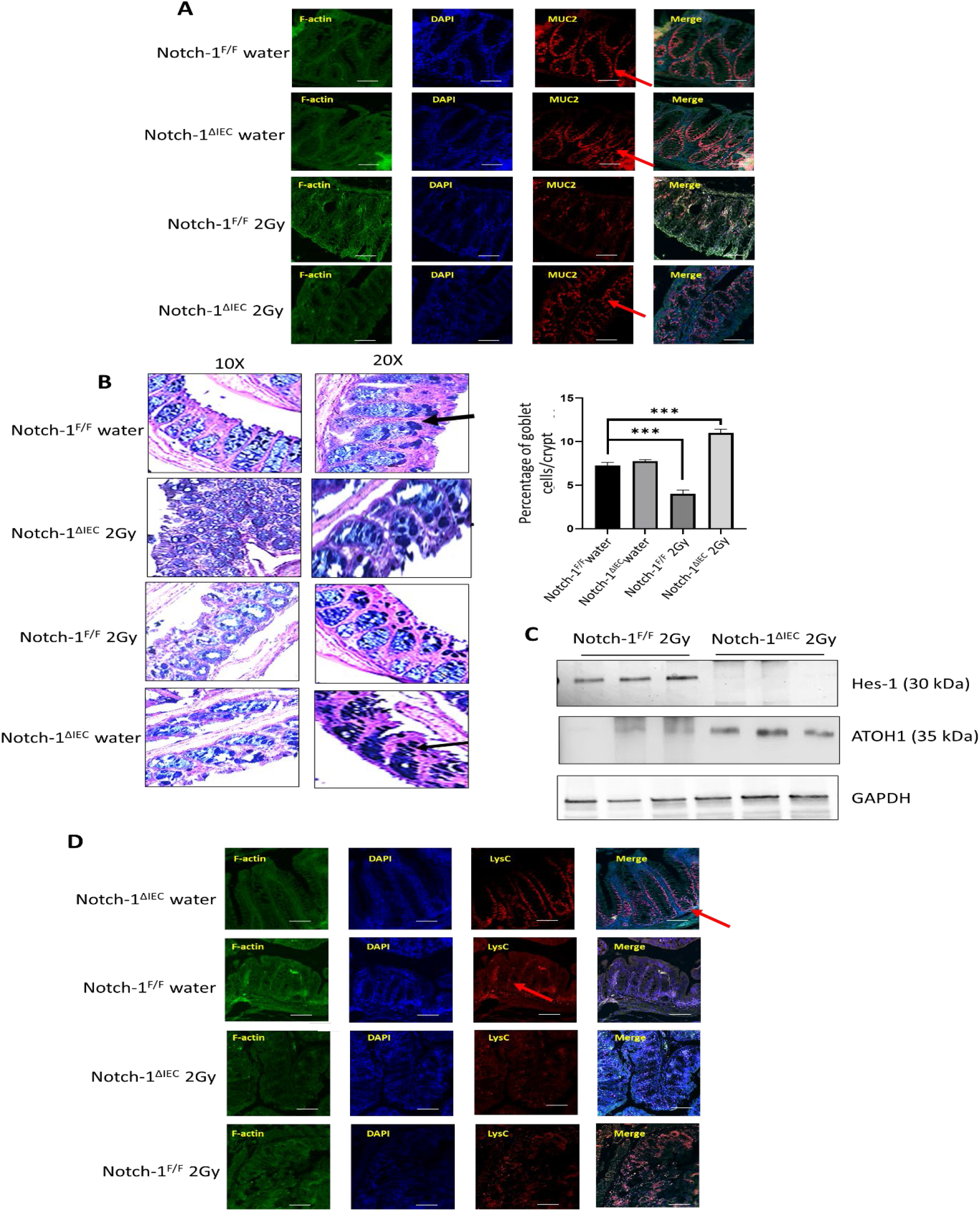
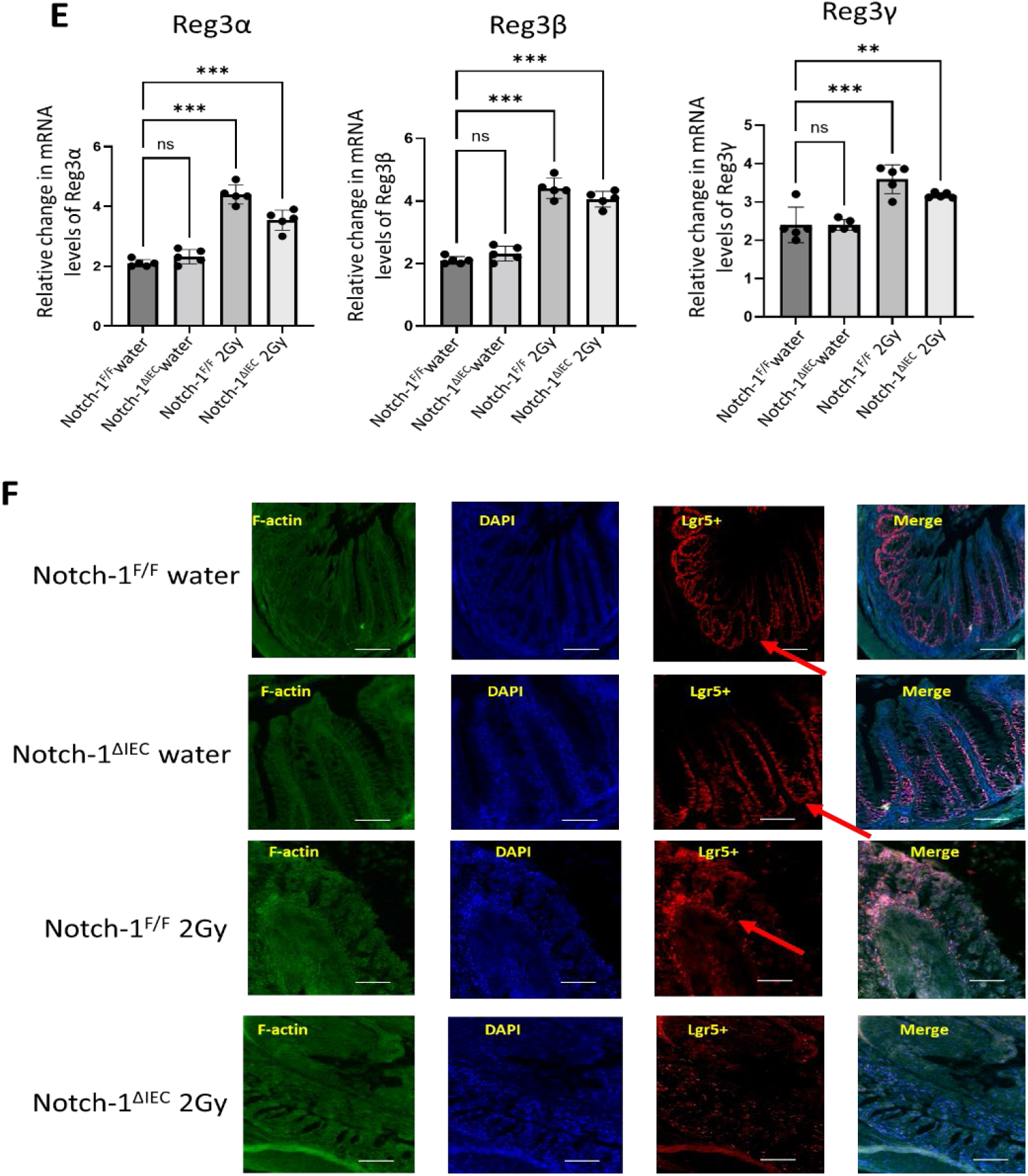
Notch-1 modulates the expression of antimicrobial proteins and inhibits secretory cell lineage differentiation in colonic epithelium. (A) Immunofluorescence staining for MUC2 in the colon of 8 week-old Notch-1^ΔIEC^ mice and control Notch-1^F/F^ (2Gy and water group) using mAbs against MUC2 (red), nuclei were counterstained with DAPI (blue). Scale bars, 100 μm. (B) AB-PAS staining for detection of goblet cells. The number of goblet cells per crypt was quantified. Values are mean ± SD (n = 5). ***p < 0.01 (Mann–Whitney U test). (C) Immunoblotting for Hes-1 and ATOH1 in membrane protein extracts obtained from the colon of radiation-induced colitis mice groups (Notch-1^ΔIEC^ and Notch-1^F/F^). (D) Immunofluorescence staining for Lysozyme C in the colon of 8-week-old Notch-1^ΔIEC^ mice and control Notch-1^F/F^ (2Gy and water group) using mAbs against Lysozyme C (red), nuclei were counterstained with DAPI (blue). Scale bars, 100 μm. (E) qRT-PCR was used to examine the mRNA expression of Reg3α, Reg3β, and Reg3γ. Data were expressed as mean ± SD of three independent experiments. (0ne-way ANOVA) * p < 0.05, ** p < 0.01, *** p < 0.001, **** p < 0.0001, vs. radiation groups (Notch-1^ΔIEC^ and Notch-1^F/F^). (F) Immunofluorescence staining for Lgr5+ in the colon of 8 wk-old Notch-1^ΔIEC^ mice and control Notch-1^F/F^ (2Gy and water group) using mAbs against Lgr5+ (red), nuclei were counterstained with DAPI (blue). Scale bars, 100 μm.

Notch-1 deletion activated secretory cell lineage differentiation. Western blotting data shows that Notch-1 mediates Hes-1 and inhibits ATOH1 expression and hence suppresses goblet cell proliferation (Fig 7C). Enhanced expression of lysozyme showed that the Paneth cell population was expanded in Notch-1^ΔIEC^ mice as compared to Notch-1^F/F^ mice after radiation-induced colitis[28] (Fig 7D). Immunostaining of lysozyme identified Paneth-like cells in the colonic epithelium of Notch-1^ΔIEC^ and Notch-1^F/F^ mice[29]. We subsequently examined the expression of antimicrobial proteins in the colonic epithelium (Fig 7E). The expression of the regenerating islet-derived (Reg) gene family encoding Reg3α, Reg3β, and Reg3γ in the colonic epithelium of Notch-1^ΔIEC^ and Notch-1^F/F^ mice in different conditions. Interestingly, The expression of antimicrobial proteins Reg3α, Reg3β, and Reg3γ, all of which are known to be secreted by Paneth cells [30, 31], was remarkably upregulated in the colonic epithelium of Notch-1^ΔIEC^ mice without any significant difference from Notch-1^F/F^ mice after radiation insult. Collectively, ablation of Notch-1^F/F^ unlikely attenuates the production of antimicrobial proteins by IECs. Radiation depletes Lgr5+ cells [32] [33]. Notch-1 overexpression in the crypt after ablation of Lgr5+ regenerates stem cells [33]. (Fig 7F) Lgr5+ cells were seriously downregulated after radiation exposure in Notch-1^ΔIEC^ mice. However, some regeneration was seen in Notch-1^F/F^ mice even after the insult. This shows that Notch-1 mediates regeneration and proliferation in the colon crypt.

### Notch-1 is Essential for Injury-Induced Intestinal Stem Cell Recovery Through its Ligands-Controlled Expression Under the Switch of the KLF5 Modulator

Intestinal Stem Cells (ISCs) replenish the epithelium daily. However, radiation-induced colitis ablates Lgr5 ISCs. In return, this was found to activate stemness’s function in slow-cycling progenitors or even differentiated epithelial cells, most notably secretory lineages [34]. Notch signaling pathway is required for ISCs to maintain stemness. KLF5 is an epithelial cell-intrinsic regulator essential for epithelial regeneration after injury-induced ISC attrition. Klf5 deacetylation activates NOTCH signaling, Hes1, Jagged 1, and DLL-1, which promotes luminal cell proliferation [35, 36]. To investigate if Notch-1 affects KLF5 directly, we performed immunostaining using KLF5 specific antibody in Notch-1^ΔIEC,^ and Notch-1^F/F^ colon mucosal epithelial cells in radiation caused colitis condition vs. normal no radiation (Fig 8A). Notch-1^F/F^ mice significantly lost KLF5 expression post-radiation. However, the ligand responsible for such an increase in Notch activation remains uncertain. Therefore, we examined the expression of DLL-1, DLL-3, DLL-4, JAG-1, and JAG-2 in colitic mucosal tissues of radiation-colitis mice. (Fig 8B) RT-qPCR analysis showed a surprising loss of Dll1 expression in Notch-1^F/F^ radiation-colitis mice. Conversely, a striking increase in the mRNA expression of Dll4 was observed in Notch-1^F/F^ 2Gy, suggesting that a distinct regulation of Dll1 and Dll4 expression exists under the inflammatory environment. (Fig 8C) DLL-3 and JAG-2 didn’t show any significant difference from the control, but JAG-1 expression was reduced in Notch-1^F/F^ and Notch-1^ΔIEC^ mice post-radiation. Dll4 and JAG-1 may act as a significant Notch ligand in the crypts of the inflamed colonic mucosa. It supported the conclusion that Notch-1 lowers the expression of KLF5 and hence mucus producing cells known to proliferate in the presence of KLF5. Notch-1 doesn’t let the cells differentiate in the transit-amplifying region; rather dedifferentiates them for ISC renewal. As the secretory cells decrease during colitis, DLL-1 primary ligand expressed by goblet cells also decreases.

**Fig 8.**
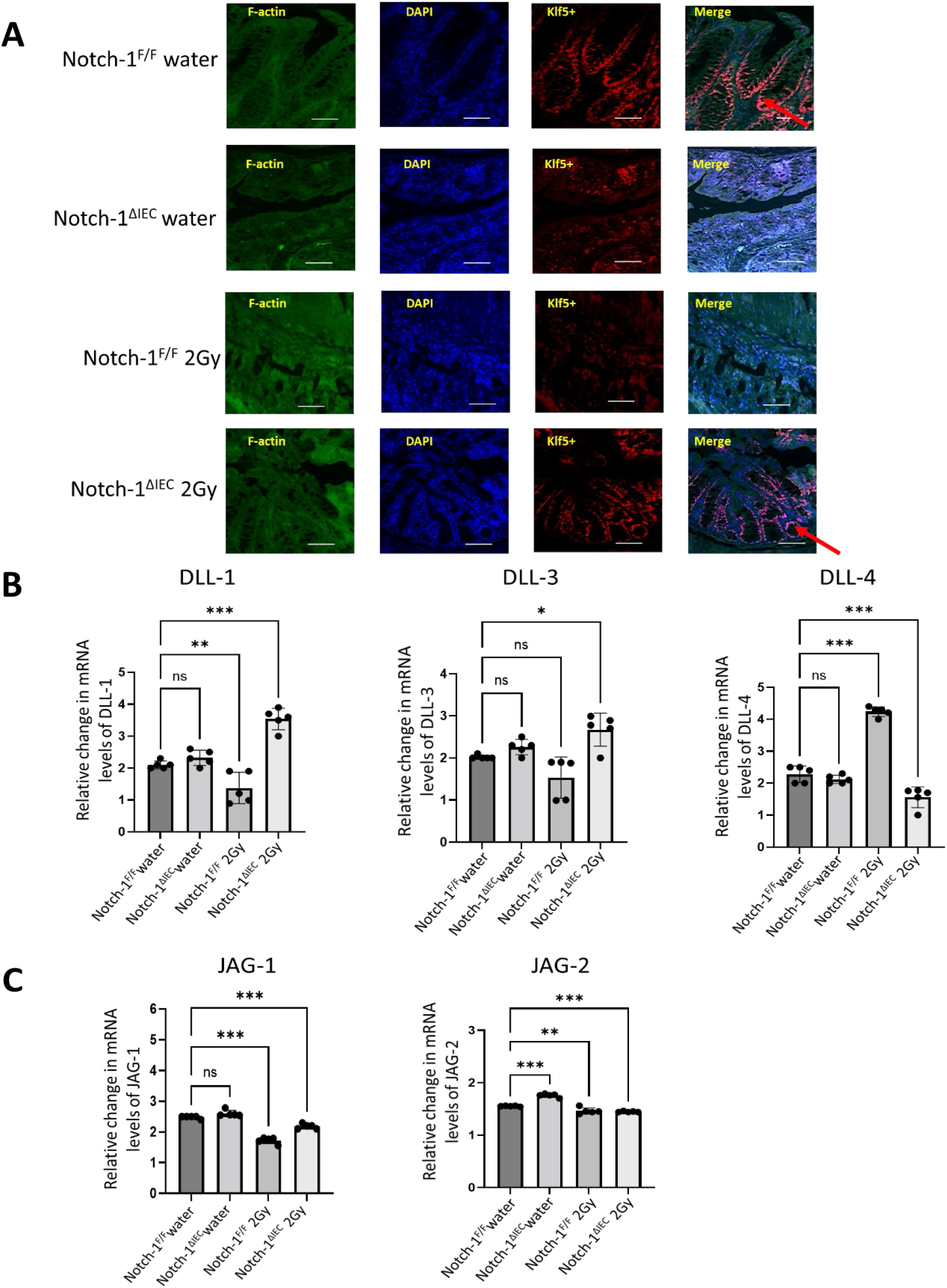
Role of Notch-1 in injury-induced healing. (A) Colonic tissue sections of 8 wk-old Notch-1^ΔIEC^ mice and control Notch-1^F/F^ (2Gy and water group) were stained with anti-KLF5 antibody (red), nuclei were counterstained with DAPI (blue). Scale bars, 100 μm. (B) Expression of Notch ligands in the intestinal epithelium of Notch-1^ΔIEC^ and Notch-1^F/F^ littermates, RT-qPCR was used to examine the mRNA expression of DLL-1, DLL-3, and DLL-4 (C) The expression of more Notch ligands JAG-1 and JAG-2 were analyzed by RT-qPCR in colonic epithelial mucosa. Data were normalized to the expression of 36BR. Data were expressed as mean ± SD of three independent experiments (n=5) * p < 0.05, ** p < 0.01, *** p < 0.001 (one-way ANOVA).

### Notch-1 Regulates Mucosal Inflammation Through Interaction with Cytokines

Radiation-induced colitis severity was assessed by colonoscopy and colonic levels of TNF-α, IL-6, IL-12, IL-22, and IL-10. The widely used murine colitis model, IL10^-/-^, has a normal inner mucus layer in terms of thickness but produces a mucus structure that is penetrable to bacteria [37]. It suggests the role of cytokines in mediating the intestinal barrier. To decipher how Notch-1 modulates the expression of cytokines, we performed qPCR for different pro-inflammatory and anti-inflammatory cytokines. Compared with the control Notch-1^F/F^ H_2_O group, the mRNA levels of IL-6, IL-22, IL-12, TNFα, and IL-10 were significantly elevated in colonic tissues of mice with radiation-induced colitis (Fig 9A). According to previous research, Notch signaling is also necessary to produce IL-22 [38]. Interestingly, the expression of IL-10, an anti-inflammatory cytokine, also increased in mRNA QPCR experiments following the radiation induction on day 7. The concentration of pro-inflammatory cytokines mentioned above was drastically increased in the Notch-1^F/F^ 2Gy group compared to the Notch-1^ΔIEC^ 2Gy group. This confirms that Notch-1 spikes the pro-inflammatory cytokines.

**Fig 9.**
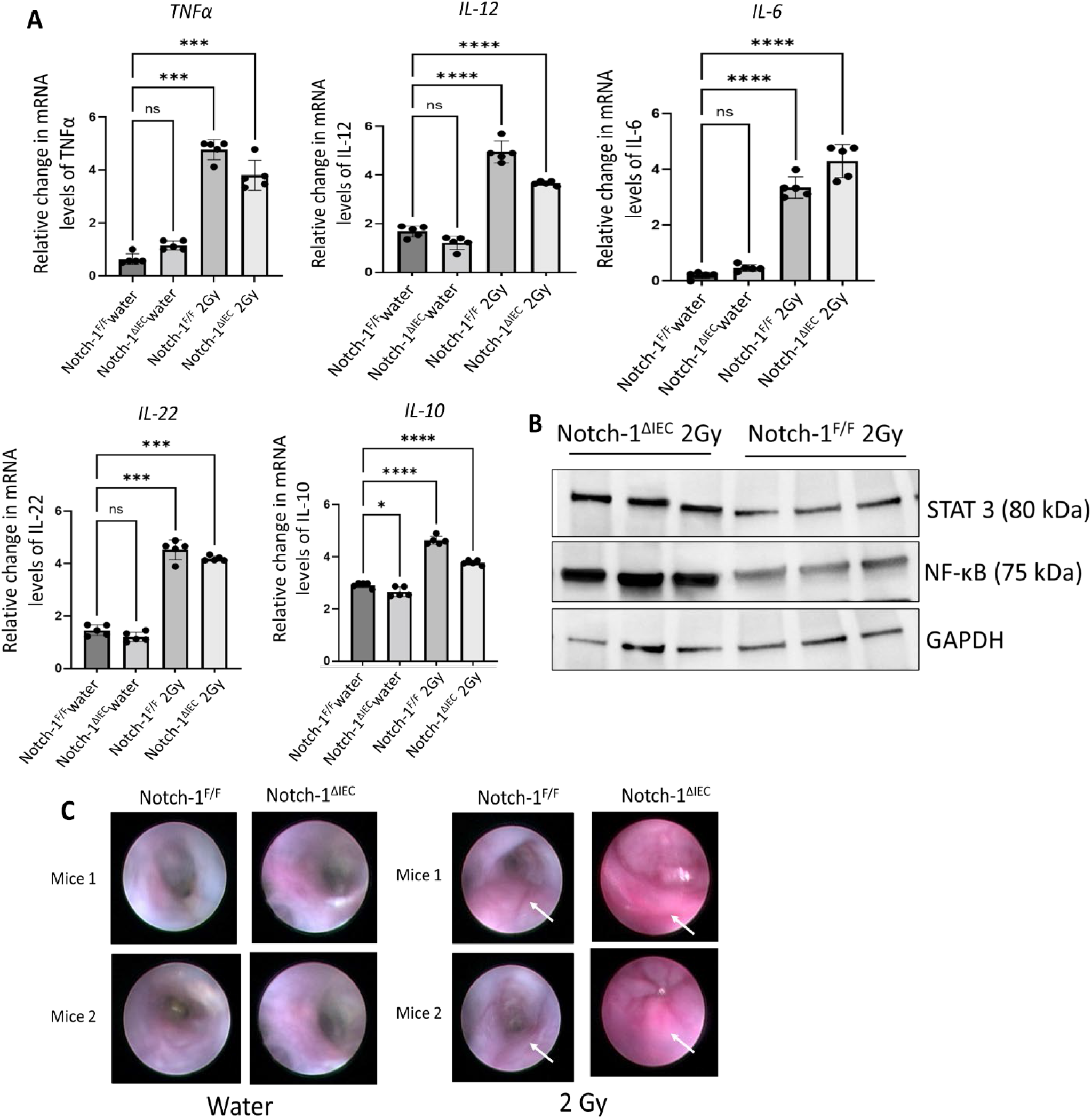
Notch-1 affects pro-inflammatory and anti-inflammatory cytokines production in mice with radiation-induced colitis. (A) mRNA expression of cytokines including TNF, IL-6, IL-12, IL-22, and IL-10 was examined by qRT-PCR in colonic mucosa from 8-week-old mice after seven days of recovery from radiation injury. Data is depicted as fold change. Data were normalized to the expression of 36BR (housekeeping gene). Data were expressed as mean ± SD (n=5) of three independent experiments. * p < 0.05, ** p < 0.01, *** p < 0.001, **** p < 0.0001, vs. radiation groups (one-way ANOVA test followed by Dunnett test). (B) Colon of mice (Notch-1^ΔIEC^ and Notch-1^F/F^ either exposed to 2Gy radiation or not (control)) was extracted and longitudinal dissected. Mucosal scrapings were taken to analyze the protein level of Nf-kB and STAT 3 to find their association with Notch-1. Immunoblot for the expression of STAT3 with GAPDH as a loading control shows downregulation of STAT3 during the upregulation of Notch-1 in Notch-1^F/F^ mice. (C) Colonoscopy images to show inflammation in colon increased slightly in Notch-1^F/F^ mice after radiation induced colitis but Notch-1^ΔIEC^ mice had escalated inflammatory response. The area focused in white squares shows shows propensity of inflammation in different groups. Scale bars, 100 μm.

Notch/STAT3 is a universal molecular switch regulating anti-inflammatory functions in chronic inflammation [19]. Notch signaling integrates STAT3-dependent inflammatory cytokine cues to induce IL-10 expression selectively. To test this, we checked immunoblotting data, which clearly shows the tapering of STAT 3 protein in the presence of Notch-1, which results in less IL-10 production in the presence of Notch-1 in acute colitis (Fig 9B).

To investigate the effect of radiation exposure on the colon of Notch-1^ΔIEC^ and Notch-1^F/F^ mice, we visualized mucosal inflammation by colonoscopy. Inflammation in the colon for different groups showed a similar tendency depicted by the rise in cytokines. (Fig 9C). Thus, our data confirm that inhibiting the expression of Notch-1 in the later stages of UC could inhibit the overwhelming inflammatory response, which is partly responsible for the treatment of UC.

### Gut Microbiota Population is Altered in Conditional Notch-1 Deficient Colon

Notch-1 has a trophic effect on the epithelial barrier, contributing to their direct inhibitory effect on bacterial invasiveness. In previous studies, mouse models with spontaneous colitis have an inner mucus layer that is penetrable to bacteria [39]. Although the cause of the dysbiosis is uncertain, we hypothesized that the enhanced mucus production in Notch-1^ΔIEC^ mice colon could result in changes in the mucus-associated flora, thereby allowing bacteria with the increased inflammatory potential to become more pronounced and contribute to the development of chronic inflammation. To investigate this, we performed mRNA analysis for the 16S gene, which depicted an expected reduction in bacterial species in the mucosal scrapings of colon radiation-induced colitis mice on Day 7 post-radiation. *Bacteroidetes, Fusobacterium*, and *Enterococcus faecalis* abundance increased significantly following radiation injury in Notch-1^F/F^ mice (Fig 10A). However, the most abundant mucolytic genera, *Akkermansia muciniphila*, remained higher in the Notch-1^ΔIEC^ irradiated group. *Firmicutes* abundance plummeted after radiation in both Notch-1^F/F^ and Notch-1^ΔIEC^ mice. RT-qPCR also quantified Muc2 gene expression in all the groups of Notch-1^F/F^ and Notch-1^ΔIEC^ mice (with/ without radiation), confirming the trends formerly pronounced, statistically significant in RT-qPCR data of different bacterial phyla (Fig 10B). In sum, these data suggest that disruption in the luminal environment of Notch-1 deficient mice contributes to an alteration in the composition of bacterial flora associated with the colonic mucosa, likely initiating the pronounced inflammation.

**Fig 10.**
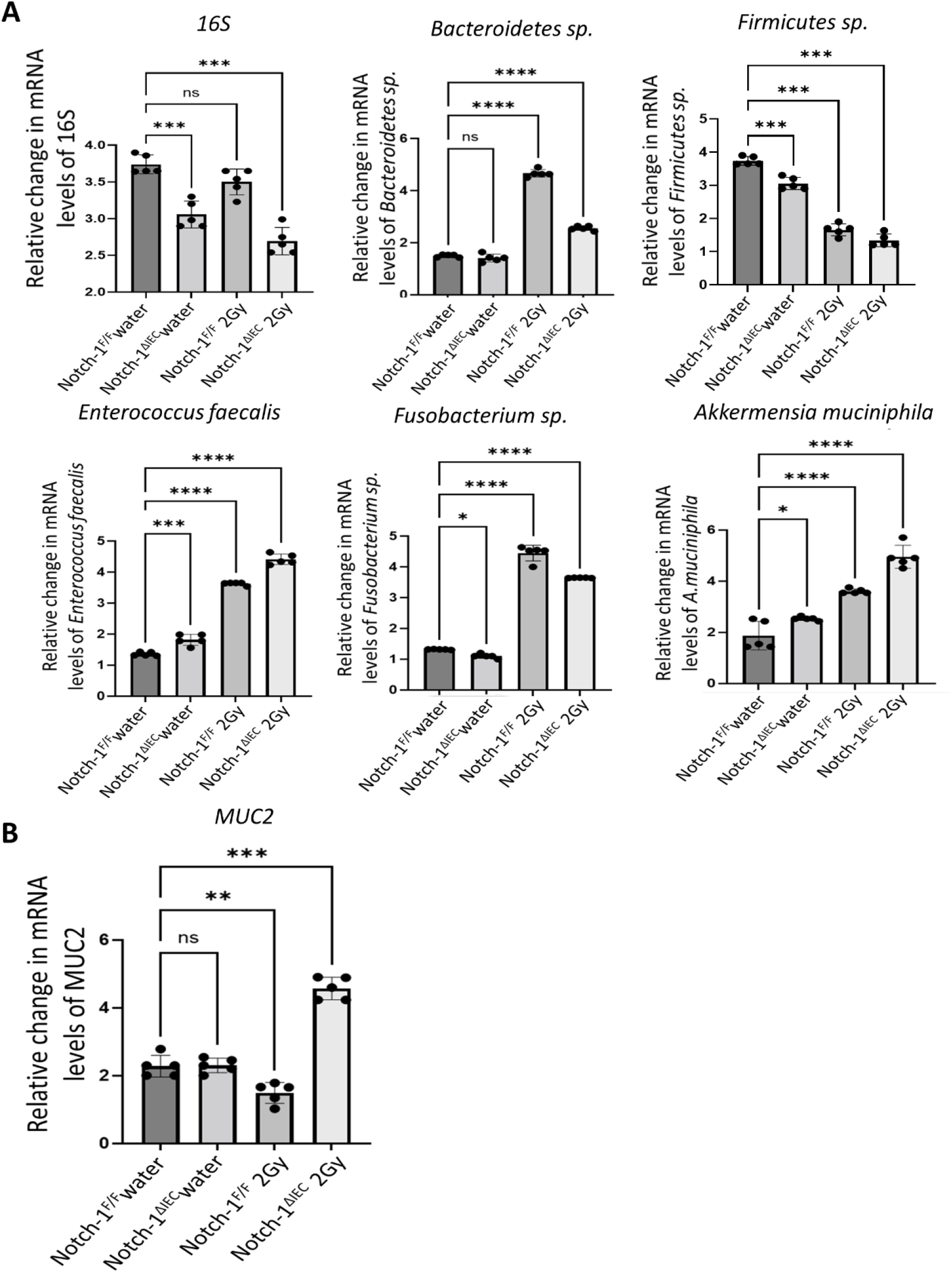
Notch-1 affects gut microbiota composition. (A) mRNA expression of different commensal bacteria by RT-qPCR, including total bacterial population 16SrRNA, Firmicutes, Bacteroidetes sp., Fusobacterium, Akkermensia (A. muciniphila), and Enterococcus faecalis (E. faecalis) was examined by qRT-PCR in colonic mucosa from 8-week old mice after seven day recovery from radiation injury. (B) RT-qPCR for MUC2 gene. Data is depicted as fold change. Data were normalized to the expression of 36BR (housekeeping gene). Data were expressed as mean ± SD (n=5) of three independent experiments. * p < 0.05, ** p < 0.01, *** p < 0.001, **** p < 0.0001, vs. radiation groups (one-way ANOVA test followed by Dunnett test).

**Fig 11.**
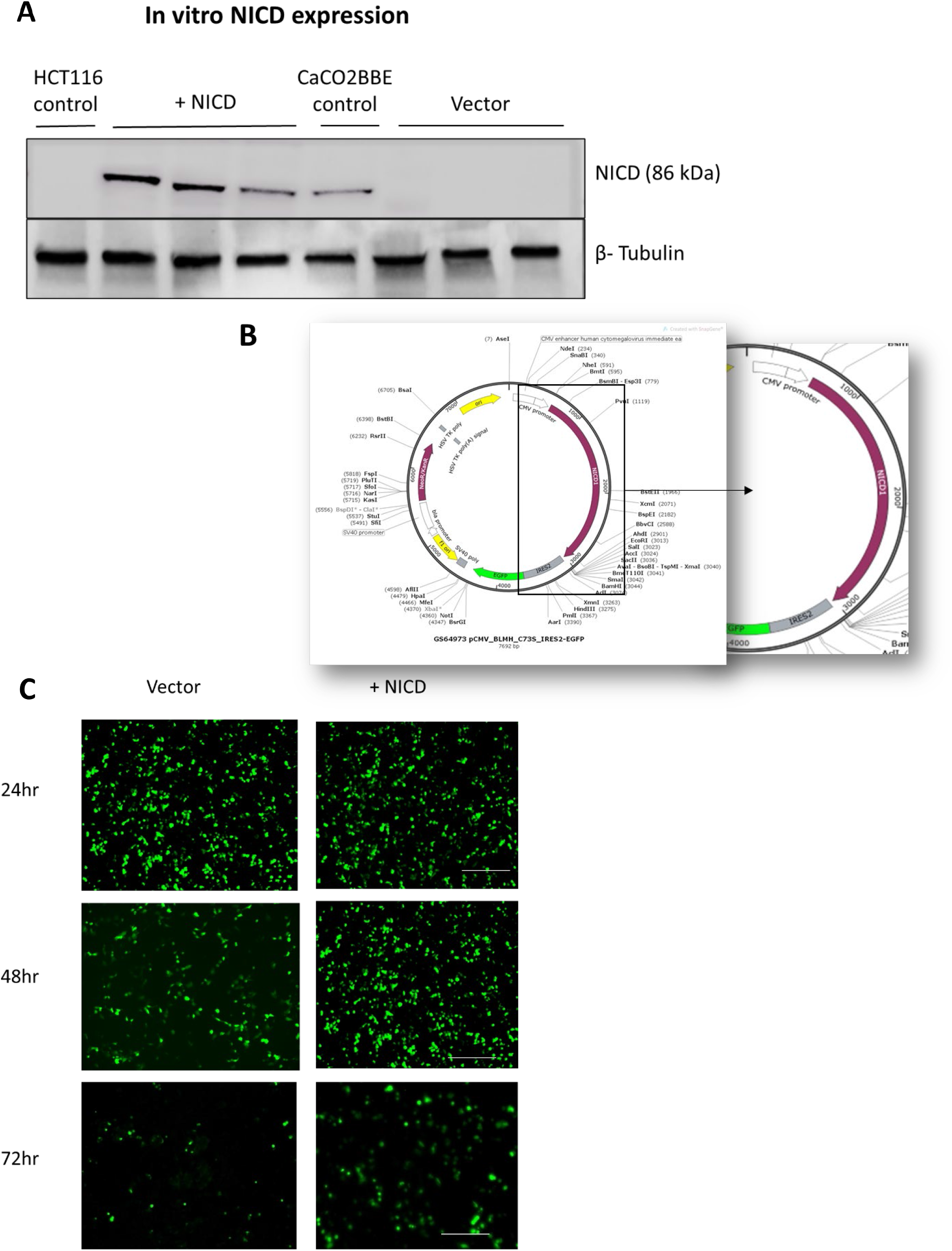
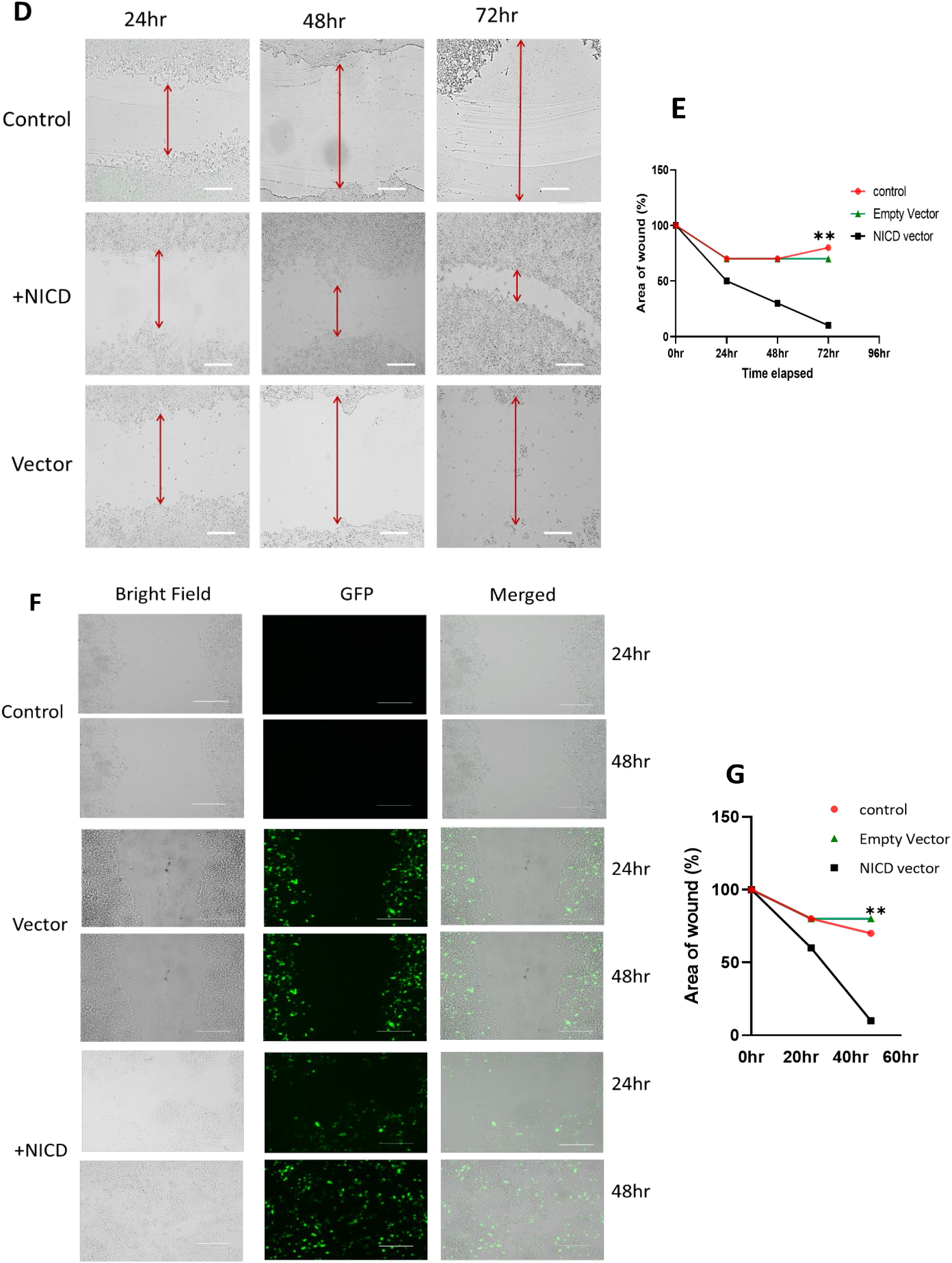
In vitro wound healing assay to access Notch-1 role in wound recovery in colitis. (A) Immunoblotting of the cells (HCT116 and Caco-2) as controls and HCT116 transfected by NICD1 or empty plasmid using Ab against NICD1. Data are representative of three independent experiments. (B) Illustration of plasmid used in transfection experiments(C) Transfection efficiency showed via EGFP expression in fluorescence microscope at different time intervals (24, 48, and 72 hr). Scale bars, 100 μm. (D) Representative images of cell filling gap via cell proliferation at 24hr, 48hr, and 72hr in control (vehicle), Empty vector, and NICD1 plasmid conditions were subjected to scratch wounding. Scale bars, 100 μm. (E) Quantitative data is shown in area of wound recovery. Percentages were expressed as mean ± SD of three independent experiments. (XY table-area under the curve) * p < 0.05, ** p < 0.01 (0hr, 24hr, 48hr and 72 hr). (F) Representative images for cell migration after transfection of control (vehicle), Empty vector, and NICD1 plasmid at 24hr and 48hr after wounding. Images of the wound area were acquired, and the number of cells per field that had migrated into the cell-free wound zone was determined for each culture. Scale bars, 100 μm. (G) Quantitative data on cell migration, using Image J software to calculate wound width in percentage expressed as mean ± SD of three independent experiments. (XY table-area under the curve) * p < 0.05, ** p < 0.01 (0hr, 24hr, 48hr and 72 hr).

### Notch Signaling Regulates the Motility and Proliferation of Intestinal Epithelial Cells

To examine if Notch-1 helps in wound healing in colitis. We performed an in vitro wound healing scratch assay. Studying the collective proliferation of cells in a two-dimensional confluent monolayer in highly controlled in vitro conditions allows us to investigate the role of NICD1 or lack thereof in wound healing. For the assay of HCT116 cells, colon human carcinoma cells were cultured in 6 well plates and incubated for 24 hrs. When cells were 70-90% confluent, cells were then transfected with hNICD1 cloned into the pCMV-IRES2-EGFP vector or without hNICD1 (Empty vector). (Fig 11B) represents the backbone of the plasmid pCMV-IRES2-EGFP. (Fig 11C) The transfection efficiency was visualized by fluorescence microscopy, detecting the GFP signal associated with the plasmid containing the EGFP encoding gene. (Fig 11A) The presence of the NICD1 gene was also portrayed by performing an immunoblotting experiment after transfection with NICD1 and Empty vector plasmids. The controls were HCT116, and Caco-2 cell lines, which describe the presence of basal level Notch-1 in Caco-2 cells as opposed to HCT116; therefore, all the expression shown in immunoblot was using the plasmids are purely because of externally provided NICD1. Post 24 hr of transiently transfecting the plasmids into HCT116 cell line using Lipofectamine transfecting reagent. A linear thin scratch “wound” (creating a gap) in a confluent cell monolayer was made using a pipette tip (Fig 11D). Subsequently, images were captured at 24, and 48 hr regular time intervals by using a fluorescence microscope. (Fig 11D) depicts the images of the scratch assay in three conditions, control had only cells and transfecting reagent, NICD1 had pCMV-IRES2-hNICD1-EGFP vector containing notch intracellular domain gene, and Empty vector pCMV-IRES2-EGFP, which had only EGFP expression without gene of interest. It was observed visually after taking pictures with a fluorescence microscope that NICD1 transfected cells were proliferating faster to close the scratch wound, which shows that Notch-1 mediates wound healing. (Fig 11E) To quantify the effect of NICD1 in accelerating wound healing via increased proliferation, we calculated the wound width area that kept decreasing with time. ImageJ software was used to calculate the proliferation efficiency. The wound area is illustrated in percentage closure which dropped the most in NICD1 transfected wells [40].

To study Notch-1 role in migration and reduce the risk of cell proliferation confounding the study of migration, a low dose of the proliferation inhibitor mitomycin C was used. 24 hr after transfection, cells were treated with mitomycin C (10 µg/ml) for 3 hours and washed three times with PBS to inhibit cell proliferation. Mitomycin C is an antitumor antibiotic that inhibits DNA synthesis [41]. Low serum concentrations in cell medium (serum starvation) are the most common method to suppress cell proliferation in wound healing assays. (Fig 11F) The cell monolayer was then subjected to a mechanical scratch-wound induced using a sterile pipette tip. Cells were cultured for an additional 24 hours in a serum-free basal medium. Cells in the injury area were visualized under a fluorescence microscope. The wound area was calculated by manually tracing the cell-free area in captured images using the ImageJ public domain software. (Fig 11G) The migration rate was expressed as the change in the wound area over time. To quantify the migration capacity in different conditions, the wound width was expressed as the percentage of area reduction or wound closure. The closure percentage will decrease as cells migrate over time.

## DISCUSSION

The colonic epithelium comprises three lineages of cells that arise from intestinal stem cells which are enterocytes, goblet cells, and Paneth like cells [42]. Recent studies have shown that various signals such as Wnt, Sonic hedgehog, bone morphogenetic protein, and Notch interact with the stem and progenitor cells of the intestinal epithelia to finely regulate the expansion and the cell fate decision of IECs for the maintenance of the intestinal epithelia [43]. In this study, we investigated the pathway by which Notch-1 could attenuate inflammation during colitis recovery phase by inducing differentiation and proliferation of colonic epithelium resulting in faster wound healing and thereby restoring the epithelial-mucosal homeostasis.

Numerous studies have confirmed that depletion of Hes1 is associated with significant increases in the secretory lineage IECs [44]. We noticed the same in immunoblot data where Hes-1 expression was diminished from Notch-1^ΔIEC^ mice. Further studies have shown that the activation of Notch promoted the proliferation of crypt progenitor cells and directed their cell fates toward absorptive but not secretory lineage cells [3, 45, 46]. Once stimulated, activation of Notch upon ligand binding promotes the expression of Hes-1 (Hairy and Enhancer-of-split-1), which then would suppress downstream ATOH1, resulting in an increase of enterocytes in parallel with restricting differentiation into goblet cells and MUC2 expression. Goblet cells are columnar epithelial cells specializing in the secretion of high-molecular-weight glycoproteins, called mucins. Mucin 2 (MUC2), synthesized by goblet cells, is the predominant structural component of the intestinal mucus layer that functions as a barrier to protect the epithelium [47, 48]. After performing AB-PAS staining, we noticed goblet cells decrease in number or are depleted in the inflamed mucosa of Notch-1^F/F^ mice after radiation due to high expression of Notch-1; on the other hand, the goblet cell population was expanded in Notch-1^ΔIEC^ mice. MUC-2 expression in qPCR data showed congruent results depicting more mucin production in Notch-1^ΔIEC^ mice after radiation-induced colitis.

Once the epithelial layer is damaged, Notch-1 is the primary protein that supports restoring the continuity and integrated structure of the epithelium [49]. Notch-1 is responsible for the rapid expansion of undifferentiated IECs. Notch-1 also improves proliferation, as seen by the upregulation of Ki67+ cells and cyclin D1expression. Cyclin D1 expression was correlated with disease activity and cell proliferation in UC cases [44]. We observed that Ki67 and cyclin D1 expression was dominant in the crypt, where Notch expression is the highest. Ki67 expression was restored in Notch-1^F/F^ 2Gy mice, exhibiting the role of Notch-1 in proliferation. We further established that during colitis, Notch-1 prefers proliferation over differentiation. This was pointed out by the reduction of KLF5, which was evident in immunofluorescence images. Genetic network analysis identified KLF5 as a critical transcription factor regulating intestinal cell differentiation [44]. *KLF5* also participates in the cell cycle, inducing the expression of several cell cycle-related genes, including cyclin D1 and cyclin B [50, 51]. These data also raise the possibility that Notch-1^ΔIEC^ mice are defective in epithelial cell turnover and activate apoptosis. We, therefore, performed TUNEL assay, which is the hallmark of apoptosis, i.e., DNA degradation. The DNA cleavage may yield DNA breaks (nicks) which can be detected by TUNEL (**T**dT-mediated d**U**TP-X **n**ick **e**nd **l**abeling). To quantify the rate of colonic epithelial apoptosis, we counted the number of apoptotic cells per crypt. Consistent with Ki67-staining data, the number of apoptotic cells was significantly increased in the colonic epithelium of Notch-1^ΔIEC^ mice. These results suggest that Notch-1 helps maintain the intestinal epithelium barrier by preventing apoptosis.

In mouse intestinal crypts, Notch signaling is an important pathway associated with stem cell self-renewal [3, 51-54]. Accordingly, the proliferative zone of intestinal crypts contains essential Notch pathway components, such as receptors NOTCH1, ligands DLL-1, DLL-4, and JAG-1, and downstream Hes1 and Hes5 [54, 55]. To explore the role of Notch-1 in renewing Lgr5+ stem cells, we performed an immunofluorescence staining which resulted in decreased expression of stem cells after injury, but Notch-1^F/F^ mice were recuperating with the loss better than Notch-1^ΔIEC^ mice. Notch signaling is a key mechanism that regulates the balance between highly proliferative and relatively quiescent stem cells and activates asymmetric division when the tissue is under stress, providing a survival strategy for maintaining homeostasis within intestinal tissue.

Thus, these studies have suggested that Notch-1 signaling functions in the intestine regulate differentiation and proliferation of IECs, contributing to the maintenance and the homeostasis of the intestinal mucosa. However, the role of Notch signaling in tissue regeneration is less understood. Damage to the intestinal epithelia is observed in various diseases, such as acute intestinal infections, radiation injuries, or idiopathic inflammatory bowel diseases [56]. It has been established that Reg 3*α* protein plays a role in tissue regeneration as a mitogenic and/or antiapoptotic factor [50, 51], and other Reg family proteins likely have similar roles in inflamed tissues [49, 57-59]. These findings strongly suggest the involvement of Reg family proteins in regenerating the inflamed colon. Here, we demonstrated that Reg 3*α*, 3*β*, and 3*γ* are expressed in colonic epithelial cells and that their gene expression correlates significantly with the degree of histological damage to colonic tissue. We observed that Reg family mRNA expression was escalated in the presence of Notch-1 in wild-type mice that were exposed to radiation. This makes us conclude an indirect link between Notch and Reg proteins. Another change observed in the intestine during such a regenerative process is the ectopic expression of antimicrobial peptides by IECs. Paneth cells usually secrete peptides such as lysozymes, which helps maintain the ideal environment for the stem and progenitor IECs. The lower expressions of these antimicrobial peptides by IECs are frequently observed in the inflamed colonic mucosa [60, 61]. Such expressions likely support the local immune system in providing an ideal environment for regenerating the damaged mucosa. When we examined lysozyme expression via immunofluorescence, it became clear that the expression of antimicrobial peptides such as lysozymes plummeted in radiation-induced injury. It had no direct relationship with the presence/absence of the Notch-1 gene.

Advancing on finding out indirect links between Notch and wound healing, we discovered that studies of the mechanism responsible for regulating the expression of *Reg* family genes have shown that cytokines and growth factors play a critical role in the upregulation of Reg proteins, which is evident in injured mucosa [61-63]. In addition, previous studies have demonstrated that several signaling pathways and cytokines cascade with the Notch pathway to mediate epithelial regeneration, such as interleukin-22 (IL-22) and tumor necrosis factor-α (TNFα) [64, 65]. After performing a qPCR for respective cytokines, we spotted a similar trend where TNFα and IL-22, a known anti-inflammatory cytokine, had been upregulated in Notch-1^F/F^ radiation-induced colitis mice. Pro-inflammatory cytokines like IL-1β, IL-6, IFN-γ, and IL-12 were upregulated in radiation injury in Notch-1ΔIEC mice compared to Notch-1^F/F^ mice. IL-10 and IL-22, known for anti-inflammatory properties, had higher gene expression in Notch-1^F/F^ mice under stress. Further supports that Notch-1 reduces inflammation and helps restore epithelial homeostasis in colon.

Moreover, it is reported that the Notch signaling pathway contributes to maintaining tight junctions and adherens junction proteins in mice. In response to inflammation, altered claudin protein profiles in the TJ are associated with perturbed paracellular movement of fluid and solutes, which is reflected in the overall change in epithelial barrier function.

Ahmed et al. experimented by infecting the colon with *Citrobacter rodentium* and the absence of the Notch pathway. This resulted in compromised tight and adherens junctions, consequently leading to increased permeability of epithelial cells during inflammation [66]. We tested the immunoblot expression of multiple TJs, including claudin-1, claudin-2, ZO1, and occludin. Replacement of barrier-forming claudin-1 with pore/channel-forming claudin-2 ultimately influenced ion and fluid movement across cells. Expression of claudin-2 increased in the Notch-1^ΔIEC^ mice after radiation. Claudin-2 is restricted to the colonic crypt in the physiological state but extends beyond crypt-base proliferative cells in case of colitis. This results in poor epithelial barrier and inflammation due to a leaky gut.

We hypothesized that the enhanced mucus production in Notch-1^ΔIEC^ mice *in the* colon could result in changes in the mucus-associated flora, thereby allowing surface-associated bacteria with the increased inflammatory potential to become permanently established and contribute to the development of chronic inflammation. Frequent crypt necrosis was observed in the colonic mucosa of Notch-1^ΔIEC^ mice, suggesting bacterial infection. To investigate this, we performed qPCR with mucosal scrapings of Notch-1^F/F^ and Notch-1^ΔIEC^ mice in different conditions (water and radiation injury). A meaningful difference was found in the bacterial colonies where *Bacteroidetes sp, Fusobacterium*, and *E. faecalis* increased significantly following radiation injury in Notch-1^F/F^ mice. Firmicutes are far more established in a healthy colon. However, the most abundant mucolytic genera, *Akkermansia* was increased in Notch-1^ΔIEC^ mice, understandably so because *A*.*muciniphila* feeds on mucus, and due to high expression of Notch-1 in Notch-1^F/F^ mice, MUC2 is conversely diminished. In sum, these data suggest that disruption in the luminal environment contributes to an alteration in the composition of bacterial flora associated with the colonic mucosa, which likely manifests inflammation. Lack of Dll1, but not loss of Dll4 or Jagged1, causes an escalation in goblet cell numbers in the colon, suggesting that Dll1 is the most essential Notch receptor ligand in the crypt [53]. We confirmed the expression of different Delta and Jag ligands in case of radiation-induced injury and discovered that DLL-1 was reduced in Notch-1, which is congruent with the fact that DLL-1 is secreted by goblet cells and in the occurrence of inflammation goblet cells become fewer in number. Whereas in contrast, DLL-4 was augmented in Notch-1^F/F^ 2Gy mice. However, there was no observable difference in the Jag-1 and Jag-2 gene expression. Suggesting DLL-4 is the primary ligand that gets upregulated in inflammation.

To bolster our in vivo results, we validated the results by executing in vitro assays. We transfected HCT116 (human colon carcinoma) cells with a plasmid containing the NICD-1 gene and compared it with a control empty plasmid. During the migration and proliferation scratch assays, it was depicted that cells with the NICD-1 gene were healing the wound much faster than controls. This further confirmed formerly positioned in vivo results.

These results suggested that the activation of Notch-1 reduced the loss of TJs and improved cell proliferation, regulated the MUC2 synthesis in epithelial cells, and regenerated Lgr5+ intestinal stem cells in injury-induced colitis. Hence Notch-1 is the primary target to be augmented for faster wound healing. It further confirms that the activation of Notch is critical for the proper regeneration program in the epithelial layer and that it helps suppress goblet cell differentiation and promote cell proliferation. Understanding the mechanisms that lead to wound healing in IBD pathogenesis is crucial. To find new therapeutic strategies to improve patient health, Notch-1 can serve as an excellent target.

